# Early loss of Scribble affects cortical development and interhemispheric connectivity resulting in psychomotor dysregulation

**DOI:** 10.1101/780130

**Authors:** Jerome Ezan, Maité M. Moreau, Tamrat M. Mamo, Miki Shimbo, Maureen Decroo, Melanie Richter, Ronan Peyroutou, Rivka Rachel, Fadel Tissir, Froylan Calderon de Anda, Nathalie Sans, Mireille Montcouquiol

## Abstract

Neurodevelopmental disorders arise from combined defects in processes including cell proliferation, differentiation, migration and commissure formation. The evolutionarily conserved tumor-suppressor protein Scribble (Scrib) serves as a nexus to transduce signals for the establishment of apicobasal and planar cell polarity during these processes. Human *SCRIB* gene mutations are associated with neural tube defects and this gene is located in the minimal critical region deleted in the rare Verheij syndrome. In this study, we generated brain-specific conditional cKO mouse mutants and assessed the impact of the *Scrib* deletion on brain morphogenesis and behavior. We showed that embryonic deletion of *Scrib* in the telencephalon leads to cortical thickness reduction (microcephaly) and alteration of interhemispheric connectivity (corpus callosum and hippocampal commissure agenesis). We correlated these phenotypes with the identification of novel roles for *Scrib*, both cell- and non-cell-autonomous, on neuronal migration and axonal guidance respectively. Finally, we show that *Scrib* cKO mice have psychomotor deficits such as locomotor activity impairment and memory alterations. Altogether, we show that *Scrib* is essential for early brain development and that the outcomes of its brain-specific disruption support a direct or indirect participation of *Scrib* to neurodevelopmental pathologies.

## Introduction

The mammalian brain, seat of cognitive and behavioral processing, is the result of numerous, complex but coordinated mechanisms of development. Patients with disruptions in fundamental processes, such as proliferation, migration, polarity, branching or synaptogenesis will typically exhibit neurodevelopmental disorders. As a result, primary microcephaly ^1^, improper cortical layering ^2^ or commissural defects such as agenesis of the corpus callosum (ACC) ^3^ are frequently associated with neurobehavioral disorders including Intellectual Disabilities, Autism Spectrum Disorders (ASDs), epilepsy and/or Attention Deficit Hyperactivity Disorders (ADHD). Rare Copy-Number Variants (CNVs) associated with these disorders are found in specific regions of the human genome (including 8q24.3) and may impact these processes ^4^. In order to understand the basis of such neurodevelopmental and neuropsychiatric disorders, it is essential to decipher the genetic, molecular and cellular mechanisms that govern brain development ^5^.

Scribble (Scrib) is a conserved scaffold protein that acts as a hub for several signaling pathways and that functions in cell polarity during development ^6,7^. In rodents, a homozygous *Scrib* mutation in a spontaneous mutant called *Circletail* (*Crc*) leads to early lethality accompanied by a severe form of neural tube defects (NTDs) ^8^. This mutant displays tissue polarity defects ^9^ that are hallmarks of planar cell polarity (PCP) signaling pathway deregulation ^10^, which can ultimately impact on the development and function of the nervous system ^11–13^. Beyond its role in epithelial cells, *Scrib* has been shown to participate in motor neuron migration ^14^ and axonal guidance in the hindbrain in zebrafish^15^ but also central nervous system myelination in rodents ^16^. Additionally, our group showed a role for *Scrib* in the brains of adult mice in fine tuning of excitatory synapses and correlated its deletion with some features of ASDs ^17–19^.

In humans, the absence or mutation of the *SCRIB* gene is linked to neurodevelopmental disease. Mutation in SCRIB is associated with NTDs (OMIM # 182940) ^20–23^, like other core PCP genes ^24,25^. NTDs represent a CNS (Central Nervous System) congenital malformations that affect 1:1000 children ^26^. Spina bifida (spinal NTD) remains the commonest congenital CNS defect and is often associated with cerebellum, corpus callosum abnormalities and hydrocephalus ^27^. As a result, patients with spina bifida face neurobehavioural alterations that include psychosocial, memory and motor defects ^28^. In addition, microdeletions in 8q24.3 (encompassing both *PUF60* and *SCRIB*) were found in children presenting microcephaly ^29^. 8q24.3 deletion syndrome, also called Verheij syndrome (VRJS, OMIM #615583), is characterized by complex features such as growth retardation, short stature, dysmorphic facial features as well as renal and cardiac defects ^29,30^. The neurological symptoms of this syndrome include delayed psychomotor development, mild intellectual disability and epilepsy that are frequently associated with neurodevelopmental defects such as microcephaly and/or ACC ^29,30^. Patients with a large deletion on chromosome 8 show microcephaly and ACC ^31^, and a more specific deletion of the 8q24.3 region is associated with ASDs or ADHD ^32^. VRJS is considered as a contiguous gene syndrome because it results from the haploinsufficiency of the *SCRIB* and *PUF60* genes, which are located within the minimal critical region ^29^. The best support for *Scrib* participating in VRJS came from morpholino-based knockdown experiments in zebrafish showing altered brain size in either *scrib* or *puf60* morphants ^29^. While, all these data suggest a role for *Scrib* in early vertebrate brain development, the potential contribution of *Scrib* to some of the observed structural and psychomotor deficits has never been assessed in mammals.

To evaluate the structural and psychomotor consequences of early *Scrib* deletion in the mammalian brain, we developed a conditional gene-targeting strategy to generate two mouse lines. The results show that early deletion of *Scrib* in the dorsal telencephalon lead to 1) reduced brain cortical size associated with cortical layering defects and 2) agenesis of the corpus callosum (CC) and the hippocampal commissure. Behavioral analysis of *Scrib* cKO animals show altered psychomotor behavior accompanied by memory deficits. Altogether, our results show that the absence of *Scrib* during early brain development lead to differential structural and behavioral deficits, often observed in spina bifida or VRJS patients, supporting a role for this gene in these pathologies.

## Results

### *Scrib* expression is consistent with a developmental function in the forebrain

We evaluated the expression profile of Scrib in the developing brain using *in situ* hybridization (ISH) and an in-house Scrib antibody for immunohistochemistry. At E16.5, Scrib ISH indicated that it was expressed throughout the dorsal forebrain and enriched in specific regions such as the upper layers of the cortical plate (CP) along the medio-lateral axis (from the cingulate to the piriform cortex), in the sub-ventricular and ventricular zones (SVZ and VZ, respectively) and the Indusium Griseum (IG) at the cortical midline **(Fig 1A-A’’)**. Immunostaining confirmed this expression pattern, with enrichment in the SVZ and VZ at cell-cell junctions in both apical (labeled by Pals1) and basal domains **(Fig 1B)**. At birth (day P0), Scrib is colocalized with RC2+ radial glial cells in the cortex **(Fig 1C)** and it is also found in axonal tracts, especially in the interhemispheric commissural fibers **(Fig 1D)**. Double staining of Scrib (green) and GFAP (glial fibrillary acidic protein, a marker of mature glial cells, red) revealed that many cells among the midline glia are labeled for both markers **(Fig 1E)**.

**Fig 1.**
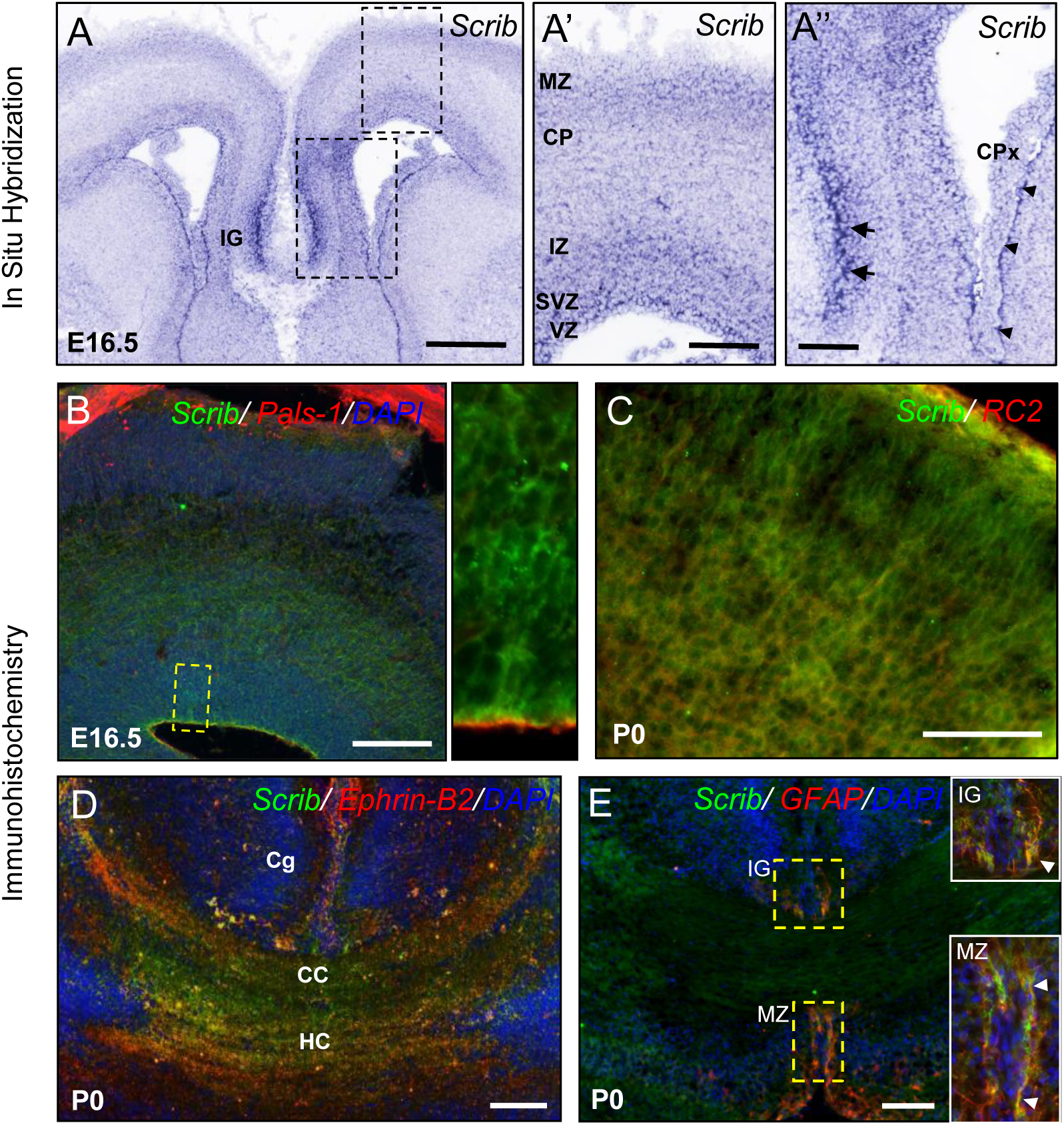
Scrib expression in the mouse developing forebrain. **A-A’’**, Representative *Scrib* expression pattern by ISH on E16.5 mouse embryo sections. The dashed box in A are magnified in A’ and A’’. Scrib mRNA is detected on the cortical plate (CP) in the subventricular zone (SVZ), the ventricular zone (VZ) and in the indusium griseum (IG, arrows) and in ependymal cells (arrowheads) next to the choroid plexus (CPx). MZ: marginal zone, IZ: intermediate zone. Scale bar: 0.5 mm (A) and 0.1 mm (A’-A’’). **B**, Representative Scrib (green) expression pattern by IF on E16.5 mouse embryo coronal sections. The dashed boxes in B are magnified in the inset. Scrib protein is detected on the cortical plate (CP) but mostly in the SVZ and the VZ. Higher magnification illustrates Scrib accumulation at cell-cell contacts in the entire VZ, but no overlap with Pals-1 (red) apical marker. Scale bar: 0.1 mm (B). **C, D, E**, Representative Scrib (green) expression pattern by IF on P0 cortex (C), Corpus Callosum (CC) (D) and midline glia (E). Scrib labeling overlaps with RC2 (marker of radial glia, red in C), Ephrin-B2 (marker of the CC, red in D) but also markedly enriched in GFAP-positive (red in E) structures including the indusium griseum (IG) and the midline zipper (MZ). Cg is the cingulate cortex. Higher magnification for selected insets (boxed areas) illustrates strong Scrib expression in glial midline structures (see arrowheads). Scale bar: 0.1 mm (E-G).

### Early *Scrib* deletion results in microcephaly

Scrib spontaneous mouse mutant *Circletail* causes severe brain and neural tube damages that result in neonatal lethality ^8^, precluding the analysis of the role of *Scrib* during forebrain development ^17^ **(Fig S1)**. In order to circumvent this issue, we have applied a conditional gene-targeting strategy to inactivate *Scrib* at different developmental stages and in different cellular types in the brain. We crossed floxed *Scrib*^*fl/fl*^ mice with either *Emx1*-Cre mice, which express the Cre recombinase starting at E10.5 in the dorsal telencephalic progenitors, or *FoxG1*-Cre mice, which express the Cre recombinase as early as E8.5 in the entire telencephalon (hereafter reported as *Emx1-Scrib*^-/-^ and *FoxG1-Scrib*^-/-^ cKOs respectively). Specific *Scrib* excision in conditional mutants was validated **(Fig 2A-F)**, and the spatial expression pattern of Cre recombinase was further confirmed by crossing *Emx1-Scrib*^-/-^ mice with the Ai6 reporter mice **(Fig 2G-H)**. The cerebral hemispheres of *Emx1-Scrib*^-/-^ cKOs were significantly smaller than those of their control littermates **(Fig 3A)**. Histological brain analysis at the rostral and caudal levels, revealed a pronounced decrease in the thickness of the caudal cerebral cortex (∼25%) **(Fig 3B)**. This decrease was observed throughout the cingulate (Cg), motor (M), primary somatosensory (S1) and secondary somatosensory (S2) cortices and was maintained in adults (data not shown). This phenotype was absent in more rostral regions of *Emx1-Scrib*^-/-^ cKO brains **(Fig 3C)**. *FoxG1-Scrib*^-/-^ mutant brains displayed a similar but more severe reduction of cortical area and thickness along the entire rostrocaudal axis **(Fig S2A-D)**. These results on cortical the cortical size reduction observed after early Scrib loss most probably arise from a combination of cell fate and migration defects.

**Fig 2.**
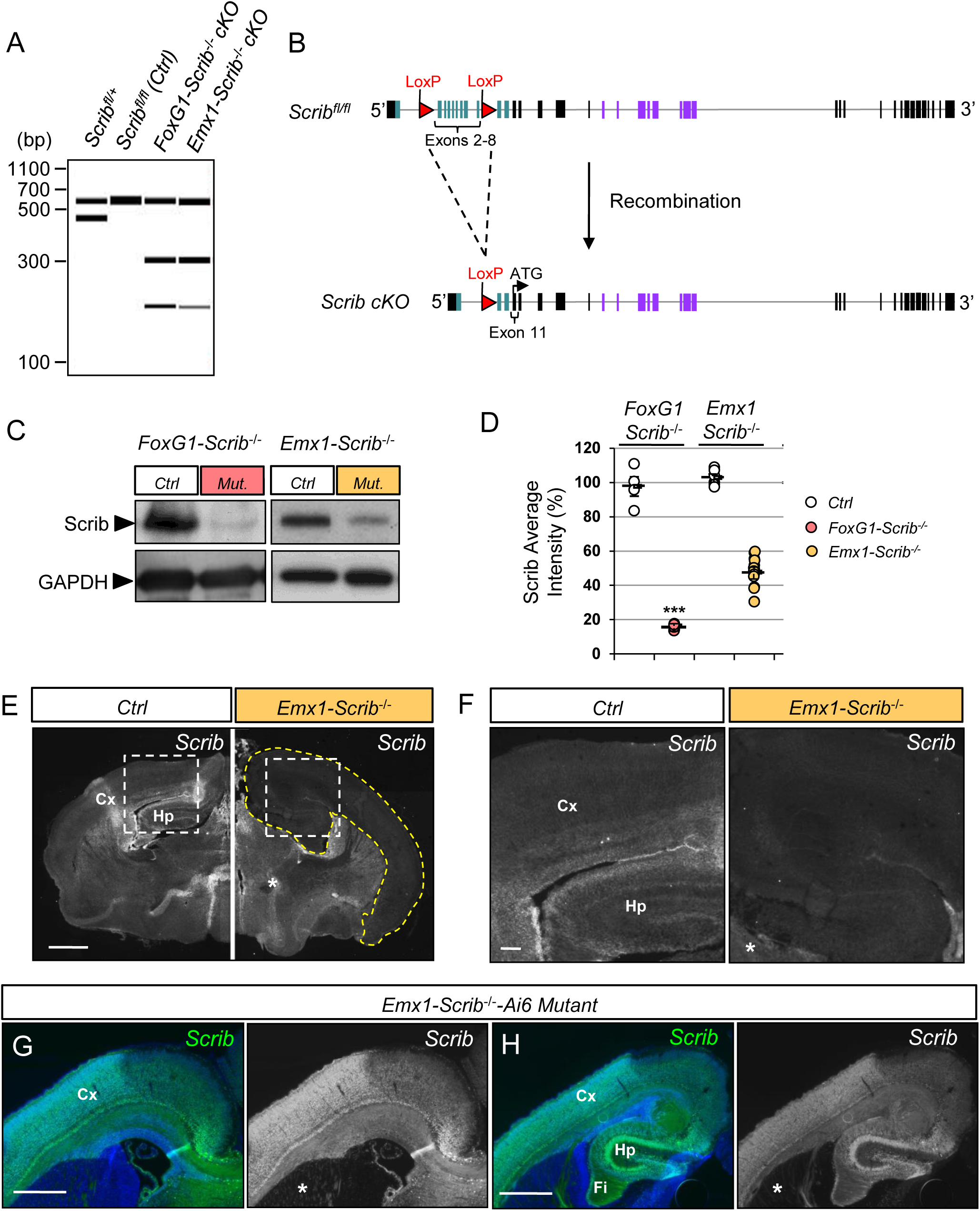
Generation and characterization of *Scrib* conditional knockout mouse mutants. **A**, PCR genotyping to detect wild-type (437 bp), floxed (541 bp) and targeted cKO (193 bp) alleles. *Cre*-mediated excision was confirmed in P0 cortices by the presence of a 300 bp product (lanes 3 and 4). In absence of Cre, the wild-type and floxed alleles remain intact as determined by 437 and 541 bp fragments, respectively (lane 1 and 2). When Cre-mediated recombination occurs (lanes 3 and 4), the floxed allele is excised, resulting in a 193 bp band, suggesting efficient recombination. **B**, Schematic representation of the genomic organization of mouse *Scrib* gene with 38 exons (solid boxes) including exons encoding for LRR (green boxes) or PDZ (purple boxes) domains of the protein. **C**, Representative western blots from control and *Scrib*^-/-^ cKO cerebral cortices showing full-length Scrib protein. Cortical protein extracts of P0 *FoxG1-Scrib*^-/-^ and *Emx1-Scrib*^-/-^ cKOs were immunoblotted with anti-Scrib antibody and anti-GAPDH as a control. cKO lysates (lanes 2 and 4) show reduced levels of Scrib when compared with control (lanes 1 and 3). **D**, Histograms summarizing western blot quantification (densitometric intensity values). Statistical analysis via a two-tailed t test (*P**<0*.*001, P***<0*.*0001*) using between 4 and 8 cortical samples per genotype from at least 3 independent experiments. Error bars indicate the SEM. **E**, Scrib expression on coronal cryosection of P0 *Emx1-Scrib*^-/-^ cKOs mutants and control littermates. A dramatic decrease in Scrib expression is observed specifically in cortex and hippocampus (area surrounded by yellow dashed line). Persistence of Scrib expression in the ventro-medial structures verifies the specificity of the excision (see star *). Scale bar: 0.5mm. **F**, Higher magnification insets for the Hp and Cx areas from E. Scale bar: 0.1mm. **G-H**, P0 coronal sections of *Emx1-Scrib*^-/-^*-Ai6* mouse brains. By crossing *Emx1-Scrib*^-/-^ with a Ai6 reporter mouse strain, we confirmed the *Emx1*-driven expression of the Cre recombinase in Cx, Hp and fimbria (Fi). Both rostral (G) and caudal (H) sections show robust ZSGreen1 expression in the dorsal forebrain assessing efficient recombination, while the striatum show almost no sign of recombination (see star *). Scale bar: 0.5mm.

**Fig 3.**
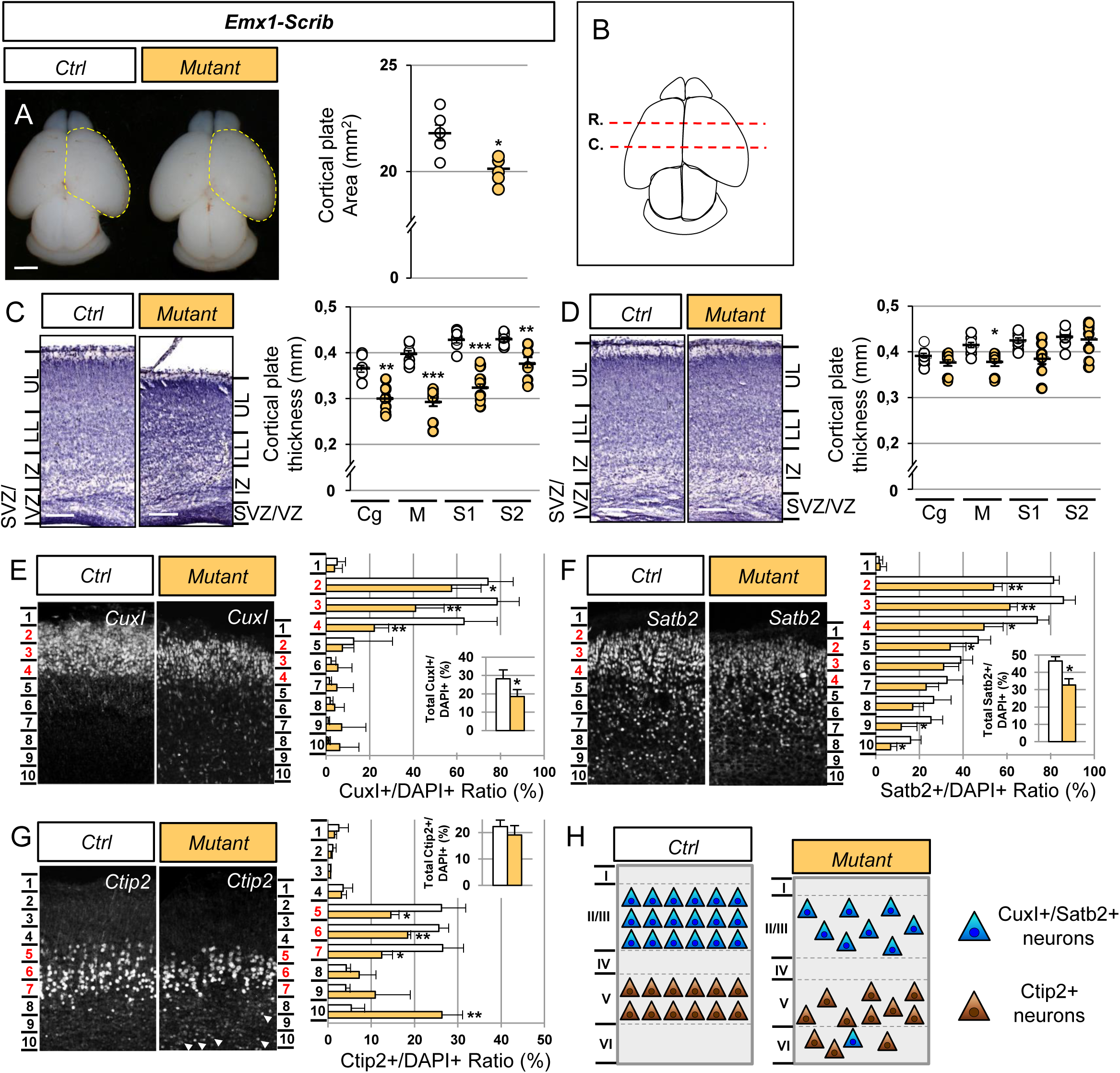
Early deletion of *Scrib* leads to microcephaly associated with cortical layering and neuronal migration defects. **A**, Dorsal views of P0 *Emx1-Scrib*-/- cKO brains. Cortical plate areas are marked with a yellow dashed line. Statistical analysis via a two-tailed t test (*P<0.05) using between 5 and 7 brains per genotype from at least 3 independent experiments. Error bars indicate the SD. Scale bar: 1 mm. **B**, Schematic view of a P0 brain sectioned coronally at the rostral (R.) or caudal (C.) level. **C-D**, Representative hematoxylin staining of coronal sections from newborn *Emx1-Scrib*-/- cKO motor cortex at the caudal (C) and rostral (D) levels and their respective controls. A marked reduction of the caudal motor cortex thickness (M) in cKOs extends to the cingulate (Cg) and somatosensory (S1-S2) cortex. No major difference was observed at the rostral level. Cortical plate thickness was measured radially from the top of the upper layer (UL) to the bottom of the lower layer (LL) of the cortex. IZ: intermediate zone, SVZ: sub-ventricular zone, VZ: ventricular zone. Statistical analysis via a two-tailed t test (**P<0.01, ***P<0.001) using 8 measurements per genotype from at least 3 independent experiments. Error bars indicate the SEM. Scale bar: 0.2 mm. **E-G**, Representative immunofluorescence staining of CuxI (E), Satb2 (F) and Ctip2 (G) on coronal sections from newborn *Emx1-Scrib*-/- cKO brains in the caudal motor cortex. Quantification of CuxI-, Satb2- and Ctip2-positive neurons is shown as a percentage (see methods). The proportion of CuxI- and Satb2-positive late-born neurons is decreased in *Emx1-Scrib*-/- cKO brains, while the Ctip2 percentage is unchanged (see insets). Several ectopic Ctip2-positive cells are mislocalized in lower bins (white arrowheads). Statistical analysis via a two-tailed t test (*P<0.05, **P<0.01) using between 3 and 4 measurements per genotype from at least 3 independent experiments. Error bars indicate the SD. **H**, Schematic representation of cortical layering in the caudal motor cortex of *Emx1-Scrib*-/- cKO and its control. Early-born neurons (brown) are mislocalized, while late-born neurons (blue) are decreased in proportion suggesting both migration and cell fate defects after early loss of Scrib function. See also Supplementary Fig S2.

### Loss of *Scrib* disrupts cortical layering

In order to further assess a role for *Scrib* in cortical layering, we next examined the expression of markers of late-born neurons, such as CuxI and Satb2 (layer II/III) and earlier-born neurons such as Ctip2 (layer V), some of these markers being essential to the formation of the CC ^33^. In *Emx1-Scrib*^-/-^ mutant P0 pups, we observed a reduction in the numbers of CuxI-positive cells (ctrl, 28.2% ± 4.8; mutant, 18.5% ± 3.7; *p* = 0.04) and Satb2-positive cells (ctrl, 46.6% ± 2.4; mutant, 32.7% ± 3.5; *p* = 0.002), suggesting defects in cell fate **(Fig 3E-F)**. This was particularly pronounced in bins 2-4, corresponding to layers II-III. In contrast, the expression of Ctip2, a marker of early-born neurons, was unchanged (ctrl, 22.3% ± 2.5; mutant, 19.1% ± 1.8; *p* = 0.24), but its cortical distribution was altered: many Ctip2-positive neurons were mislocalized in the deeper layers of the motor cortex (bins 8-10), implying migration defects **(Fig 3G)**. Notably, similar results were observed within the somatosensory cortex (data not shown). A comparative analysis in *FoxG1-Scrib*^-/-^ mutant mice showed a more severe phenotype than that of *Emx1-Scrib*^-/-^ mutants, with a dramatic reduction in CuxI, Satb2 and Ctip2 expression **(Fig S2E-H)**. These deficits probably arise from a combination of defects in cell fate and migration, as depicted in **Fig 3H**. To confirm the role of *Scrib* in neuronal migration, we used an approach based on *in utero* electroporation approach employing a previously validated shRNA construct ^17^. Scrib depletion resulted in a significant increase in the fraction of electroporated cells mislocalized in the VZ/SVZ (ctrl shRNA, 5.4% ± 0.5; Scrib shRNA, 15% ± 1.4) associated with a significant decrease in the fraction of cells reaching the upper layers (UL) (ctrl shRNA, 47.1% ± 1.4; Scrib shRNA, 30.7% ± 1.9) **(Fig 4)**. Altogether, these results indicate that embryonic deletion of *Scrib* reduces the size of the cortex, and alters the generation and positioning of the cortical neurons, including callosal projection neurons (CPNs) in the upper layers of the cortex.

**Fig 4.**
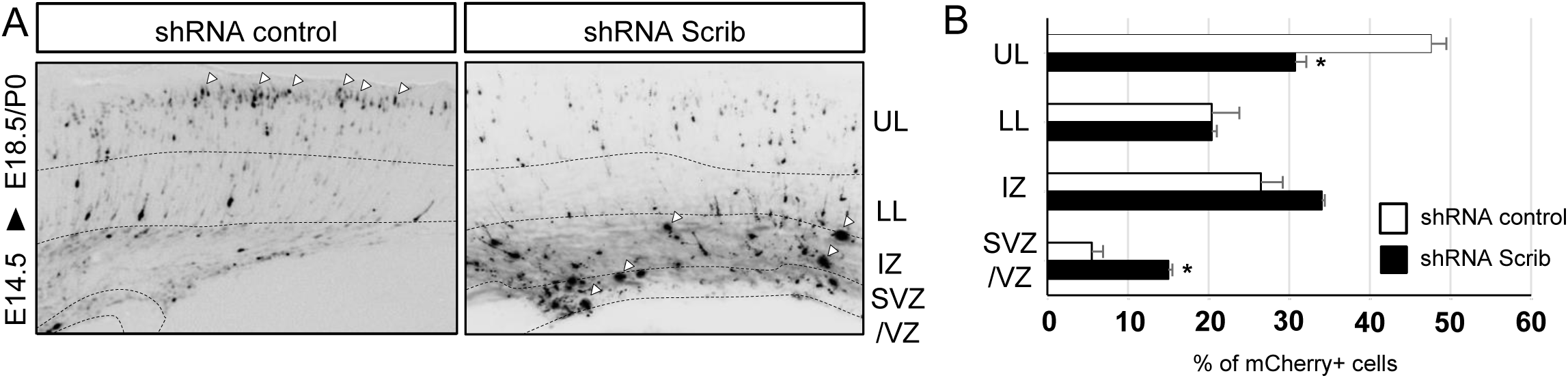
*Scrib* controls cortical neuronal migration. **A**, Cortical neurons were electroporated *in utero* at E14.5 with an mCherry expressing vector together with control shRNA or a previously validated Scrib shRNA. Brains were fixed at E18.5/P0. Nuclei were stained with DAPI (not shown) in order to delineate cortical subregions: dotted lines represent boundaries between the upper layers (UL) and lower layers (LL) of the cortex, the intermediary zone (IZ), the subventricular zone (SVZ) and the ventricular zone (VZ). Arrowheads indicate either neurons reaching the upper layers of the cortex in the control condition or Scrib shRNA-electroporated cells that remain in the deepest layers of the cortex. **B**, Quantification of the distribution of mCherry-positive cells in distinct subregions of the cerebral cortex for each condition (shRNA control, white bar; Scrib shRNA, black bar). Analysis was performed using at least 3 independent experiments. Error bars indicate SD. Statistical analysis via a two-tailed *t*-test (*\*P<0*.*01)* per condition from at least 3 independent experiments.

### Early deletion of *Scrib* affects corpus callosum and hippocampal commissure development

Histological analysis was performed along the rostrocaudal axis in order to assess the impact of early Scrib loss on the major forebrain commissure (the CC) but also the dorsal and ventral hippocampal commissure (DHC and VHC) and the anterior commissure (AC) **(Fig 5)**. Coronal histological sections of P0 *Emx1-Scrib*^-/-^ mutant brains revealed ACC in caudal sections, with callosal axons apparently unable to cross the midline **(Fig 5A-B)**. Failure of the axons to cross was accompanied by large bundles of aberrantly projecting axons near the midline, known as Probst bundles, which are frequently associated with hemispheric fusion defects **(Fig 5A’-B’)**. Anterograde axonal tracing studies using DiI staining confirmed the absence of midline-crossing callosal axons **(Fig 5A’’-B’’)**. In contrast, callosal fibers in rostral sections from the brains of *Emx1-Scrib*^-/-^ mice crossed the midline despite an apparent CC hypoplasia **(Fig 5C-D’’)**. Of note, a similar phenotype was observed in adult *Emx1-Scrib*^-/-^ mutant mice using 3D light-sheet microscopy of uDISCO treated brains injected in the caudal cortex with an adenovirus encoding GFP **(Fig 5E-F’)** suggesting that ACC is not due to a developmental delay. The deletion of *Scrib* in the *FoxG1*-Cre strain led to ACC along the entire rostrocaudal axis **(Fig S3)**. Since *Scrib* is expressed in glial midline structures **(Fig 1E)** that are critical for promoting hemisphere fusion and allowing the CC to cross between hemispheres with the help of axonal guidance signals ^34^, we next sought to determine whether CC agenesis observed in *Emx1-Scrib*^-/-^ mutant mice results from mislocalization of glial structures. Immunostaining of P0 coronal sections for GFAP and the axonal marker L1-CAM revealed that early deletion of Scrib resulted in a severe disorganization of the midline glia, with GFAP-positive cells scattered along the dorsal part of the ventricle **(FigS4)**. Parasagittal and horizontal brain sections of the *Scrib*^-/-^ mutant confirmed the presence of callosal defects. We found that the CC length, as determined as in **Fig 6A-B** was respectively reduced by ∼40% and ∼60% in *Emx1-Scrib*^-/-^ **(Fig 6C-E)** and *FoxG1-Scrib*^-/-^ **(Fig S5A-C)** mutant mice respectively, compared with control littermates. Horizontal sections from P0 *Emx1-Scrib*^-/-^ and *FoxG1-Scrib*^-/-^ mutant mice confirmed the previously observed ACC and showed a dramatic thinning of the DHC **(Fig 6F-G’, Fig S5D-E’)**. In contrast, *FoxG1-Scrib* and *Emx1-Scrib* cKOs mutant mice had intact VHC **(Fig 6H-I’, Fig S5F-G’)** and AC (arrowheads in **Fig 6C-D, Fig S5A-B)**. 3D imaging of adult *Emx1-Scrib*^-/-^ mutant mice brains cleared with uDISCO confirmed the absence of the hippocampal commissure in adult animals **(Fig 6J-K)**. Altogether, these findings indicate that early and broad deletion of *Scrib* in the dorsal telencephalon (including the cerebral cortex and the hippocampus) is responsible for a major disruption of interhemispheric commissure formation that includes agenesis of the CC and dorsal hippocampal commissure.

**Fig 5.**
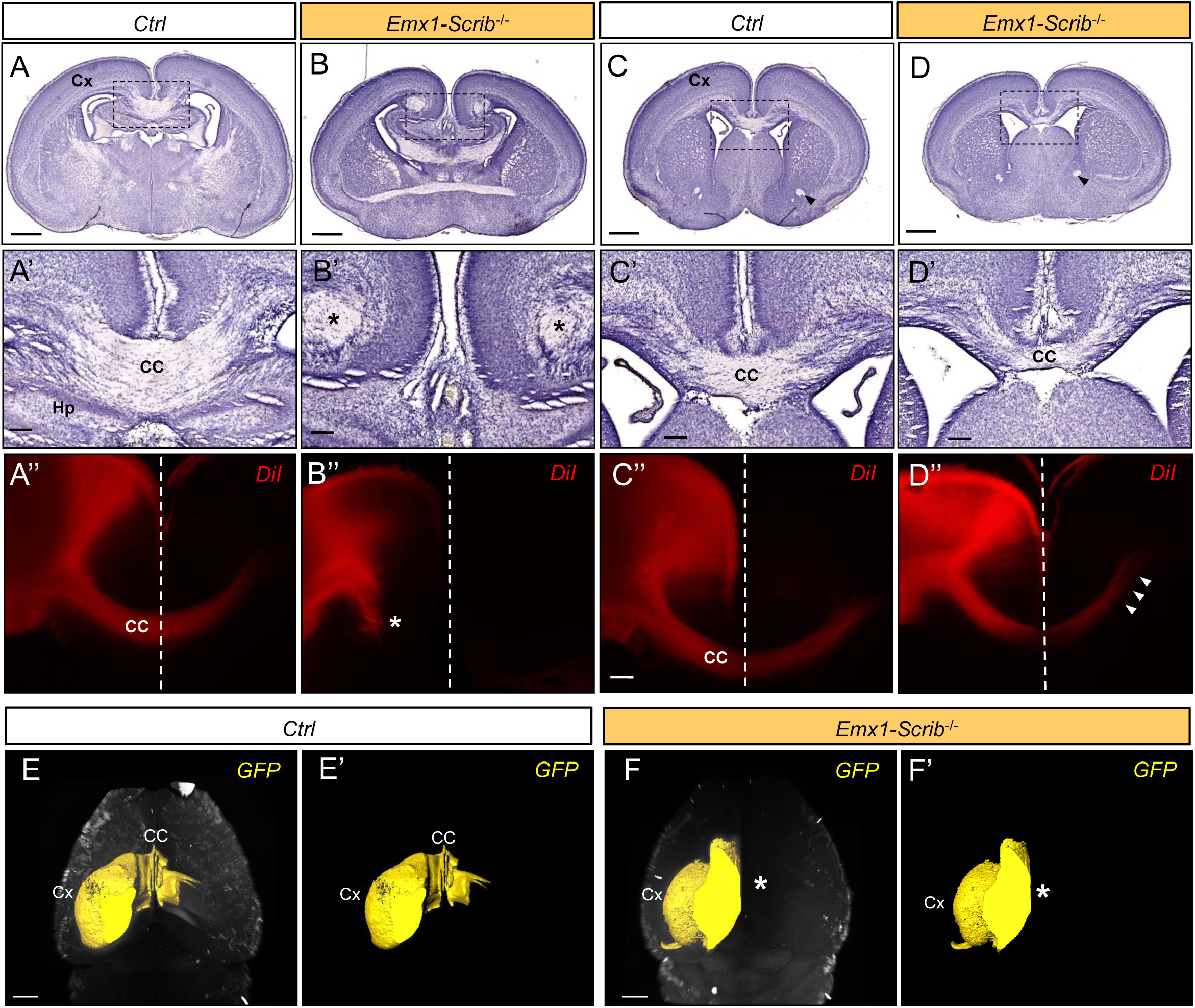
Partial corpus callosum agenesis in *Emx1-Scrib*^-/-^ *cKO* mutants. **A-D**, Representative hematoxylin staining of coronal sections from newborn *Emx1-Scrib*^-/-^ cKO brains (B and D) and their respective controls (A and C) at the caudal (A-B) or rostral (C-D) levels. Dashed boxes in A-D are magnified in A’-D’. **A’-D’**, Higher magnification for selected insets (boxed areas) from (A-D) illustrating high penetrance of CC agenesis (ACC) at the caudal level. At P0, 93% of *Emx1-Scrib*^-/-^ (n=28) cKO brains displayed ACC. Instead of crossing the midline, CC axons formed whorls (Probst bundles, PB) on either side of the midline that are indicated with an asterisk. Compared with control brains, a gap between hemispheres indicates fusion defects. Despite an apparent thinning, CC fibers do cross the midline at the rostral level. **A’’-D’’**, DiI crystals placed in the dorsomedial cortex trace CC axons in *Emx1-Scrib*^-/-^ cKO brains (B” and D”) and their respective controls (A” and C”) at the caudal (A”-B”) or rostral (C”-D”) levels at P0. *Emx1-Scrib*^-/-^ cKO^-/-^ brains displayed occasional stalled fibers in some mutant brains (asterisk in B”). Abbreviations: cortex (Cx), hippocampus (Hp), corpus callosum (CC). The midline is indicated as a white dashed line. **E-F’**, 3D imaging of adult brains cleared with uDISCO from *Emx1-Scrib*^-/-^ cKO brains (F and F’, orange) and their respective controls (E and E’). 3D reconstruction of the GFP expressed after viral infection in the sensory-motor cortex are represented in yellow. In *Emx1-Scrib*^-/-^ cKO brains, agenesis of the corpus callosum is confirmed by the absence of cortical fibers passing through the contralateral side (asterisk in F’). See also Supplementary Fig S3.

**Fig 6.**
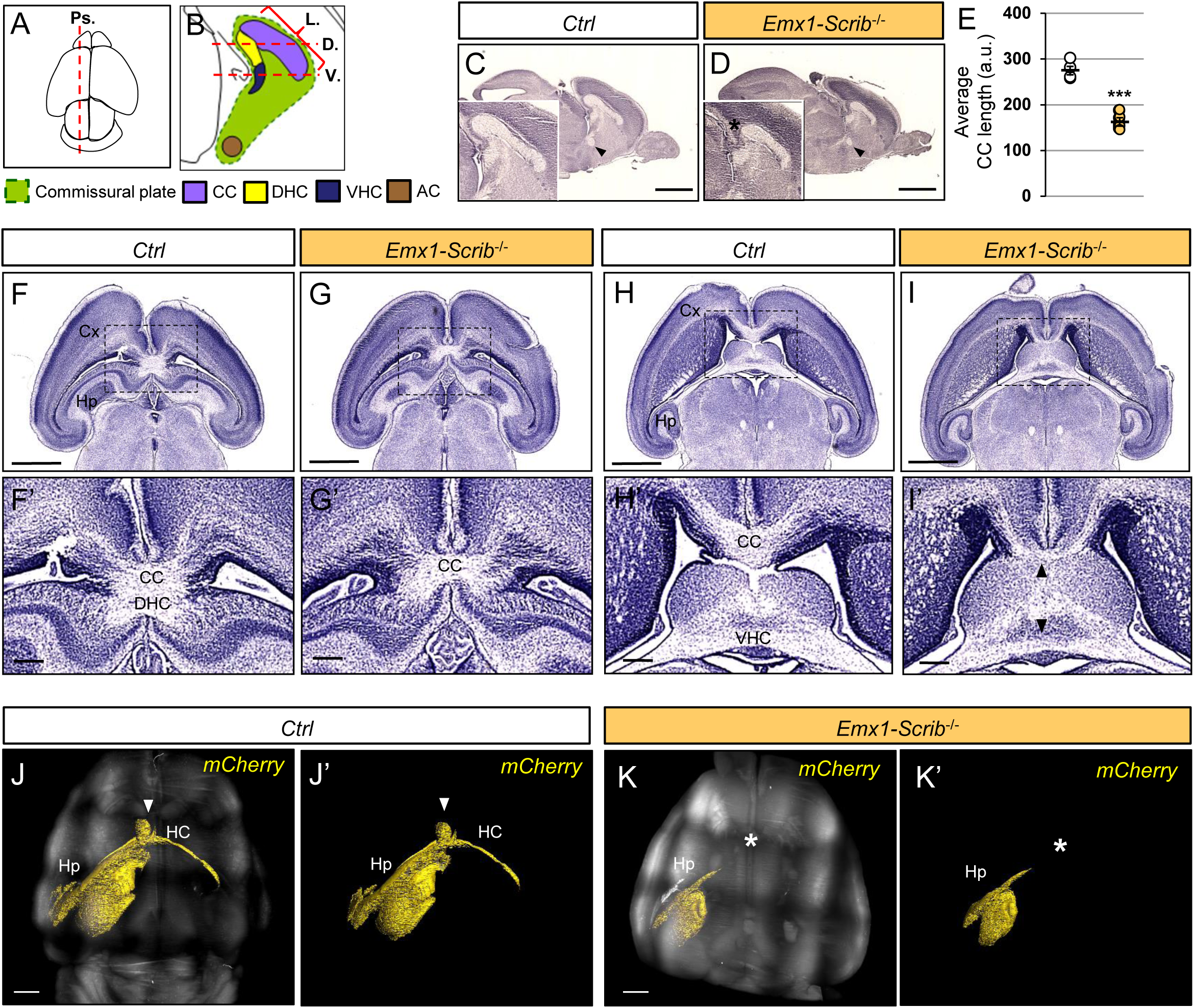
ACC is accompanied by hippocampal commissure agenesis in *Scrib*^-/-^ cKO mutants. **A-B**, Schematic dorsal (A) or para-sagittal (B) views of a P0 mouse brain. Brains were sectioned either para-sagitally (Ps. in A) or horizontally (in B) as indicated by the red dashed line. **C-D**, Marked reduction of CC length (arbitrary units, a.u.) along the rostrocaudal axis in the brains of *Emx1-Scrib*^-/-^ (D, orange, n=5) cKO brains as compared with those of their control littermates (C, white, n=5). **E**, Statistical analysis via a two-tailed *t*-test (*\*\*\*P<0*.*0001*) using between 3 and 5 brains per genotype from at least 3 independent experiments. (indicated by an asterisk). **F-I**, Representative hematoxylin staining of serial horizontal sections from newborn *Emx1-Scrib*^-/-^ (G, I) cKO brains and their respective controls (F, H) at the dorsal (F-G) and ventral level (H-I) as defined in B. **F’-I’**, Higher magnification for selected insets (boxed areas) from (F-I) illustrating agenesis of the DHC (G’) and CC hypoplasia (G’,I’). Either dorsal (F’-G’) or ventral (H’-I’) sections showed showed some callosal and hippocampal axon bundles still crossing through the midline. **J-K**, Light-sheet microscopy imaging of *Emx1-Scrib*^-/-^ cKO adult brains (K) and its control (J) cleared with uDISCO. 3D reconstruction of the mCherry expressed after viral infection in the CA3 region of the hippocampus cortex is represented in yellow. The HC (indicated by an arrowhead in control) is absent in brains from cKO mutants (asterisk in K). Scale bars: 1 mm in (C-D’,F-K), 0.2 mm in (F’-I’). See also Supplementary Fig S5.

### Early loss of *Scrib* induces hyperlocomotion and memory defects

To determine the functional consequence of the early loss of *Scrib* at a more integrated level, we next subjected our mutant model with a variety of behavioral tests that require sensory-motor, emotional and cognitive integration. Adult *Emx1-Scrib*^-/-^ cKO and their control littermates (10–20 weeks) were submitted to tests of anxiety-like behavior, locomotor and exploratory activity. The most remarkable behavior phenotype we observed was the effect on locomotor activity. First, mice were subjected to open field test to examine the basal locomotion activity exploration in a novel environment (**Fig 7A**). Compared with their controls, *Emx1-Scrib*^-/-^ mutant mice showed significantly increased activity in the open field (*t*-test: *t*_*17*_=2.35, *P<0*.*05\**; **Fig 7A**). The time spent in the center of the arena was not different between genotypes, suggesting that the *Emx1-Scrib*^-/-^ mutant mice has comparable anxiety level to control littermates (*t*-test: *t*_*17*_=0.4595, *n*.*s*; **Fig 7A**). Anxiety-like behavior was further examined in elevated plus-maze (**Fig 7B**), where *Emx1-Scrib*^-/-^ and control mice showed comparable performance, confirming that *Emx1-Scrib*^-/-^ mice have no anxiety-related behaviors (*t*-test: *t*_*17*_=1.741, *n*.*s*; **Fig 7B**). In addition, we confirmed that the *Emx1-Scrib*^-/-^ mutant mice were significantly more active in elevated plus-maze (*t*-test: *t*_*17*_=3.02, *P<0*.*01\*\**; **Fig 7B**), in Y maze (*t*-test: *t*_*17*_=2.39, *P<0*.*05\**; **Fig 8A**), and in a new cage as assessed by actimetry during a 2 hr period (genotype effect: *F*_1,13_=8.757, *P<0*.*05*\*; **Fig 7C**). In addition, *Emx1-Scrib*^-/-^ cKOs mutant mice displayed enhanced locomotor activity in their home cages during nycthemeral activity compared to the control mice, as determined by a 24-hr continuous monitoring of locomotor activities (genotype effect: *F*_1,13_=6,802, *P*<0.05*; **Fig 7D**). Hyperactivity of *Emx1-Scrib*^-/-^ cKOs mutant mice was associated with a weight loss (*t*-test: *t*_*17*_=2.412, *P<0*.*05\**; **Fig 7E**). Notably, *Emx1-Scrib*^-/-^ mice did not display changes in balance, as tested on beam walking (**Fig 7F**) and grid handling (**Fig 7G**), as well as motor coordination as tested on the accelerating rotarod (**Fig 7H**).

**Fig 7.**
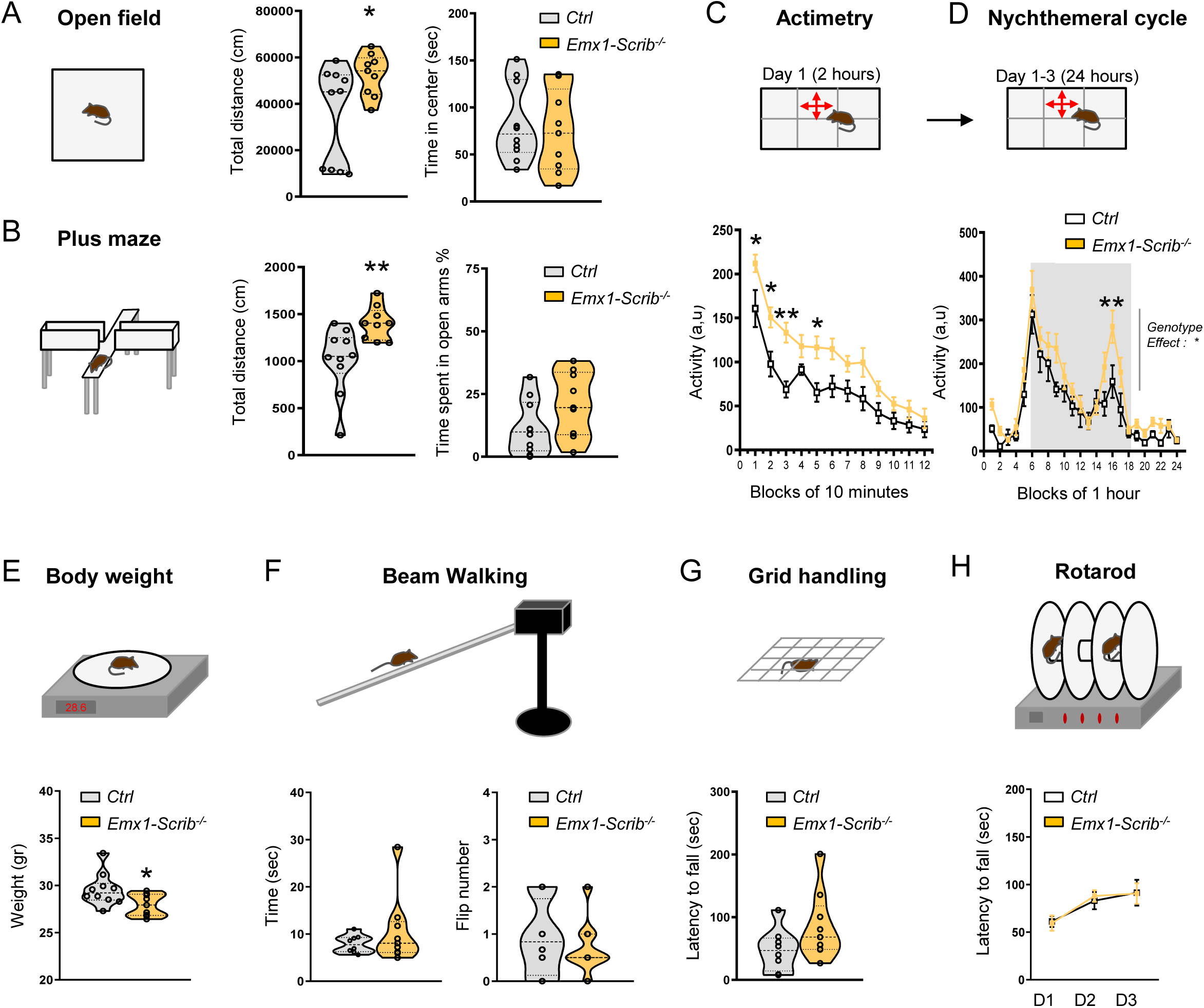
Increased activity without impaired motor coordination in Emx1-Scrib^-/-^ mice. **A**,Left panel, Open field apparatus. Middle panel, Spontaneous locomotor activity measured by the total distance in the open field test. Right panel, time spent in the center. **B**, Left panel, plus maze apparatus. Middle panel, total distance moved in the plus maze. Right panel, time spent in open arms. **C**, Upper panel, Actimetry test apparatus. Lower panel, distance travelled measured in 10-min intervals across the 120-min test session in a novel home-cage (control: n=6 mice; *Emx1-Scrib*^-/-^ : n=9 mice). **D**, Upper panel, Nychtemeral cycle test apparatus. Lower panel, distance travelled measured in 1h intervals across the 24h test session in a home cage.**E**, Upper panel, Balance apparatus. Lower panel, body weight of 10-11 week-old mice. **F**, Upper panel, Beam walking apparatus. Lower, time spent to reach the black box and total flip number. **G**, Upper panel, Grid handling apparatus. Lower panel, Latency to fall the grid. **H**, Upper panel, Rotarod test apparatus. Lower panel, Latency to fall in accelerating (4 to 40 rpm across 5 min) rotarod test on 3 consecutive days. Data are presented at means ± s.e.m. from 6-10 mice per genotype.

**Fig 8.**
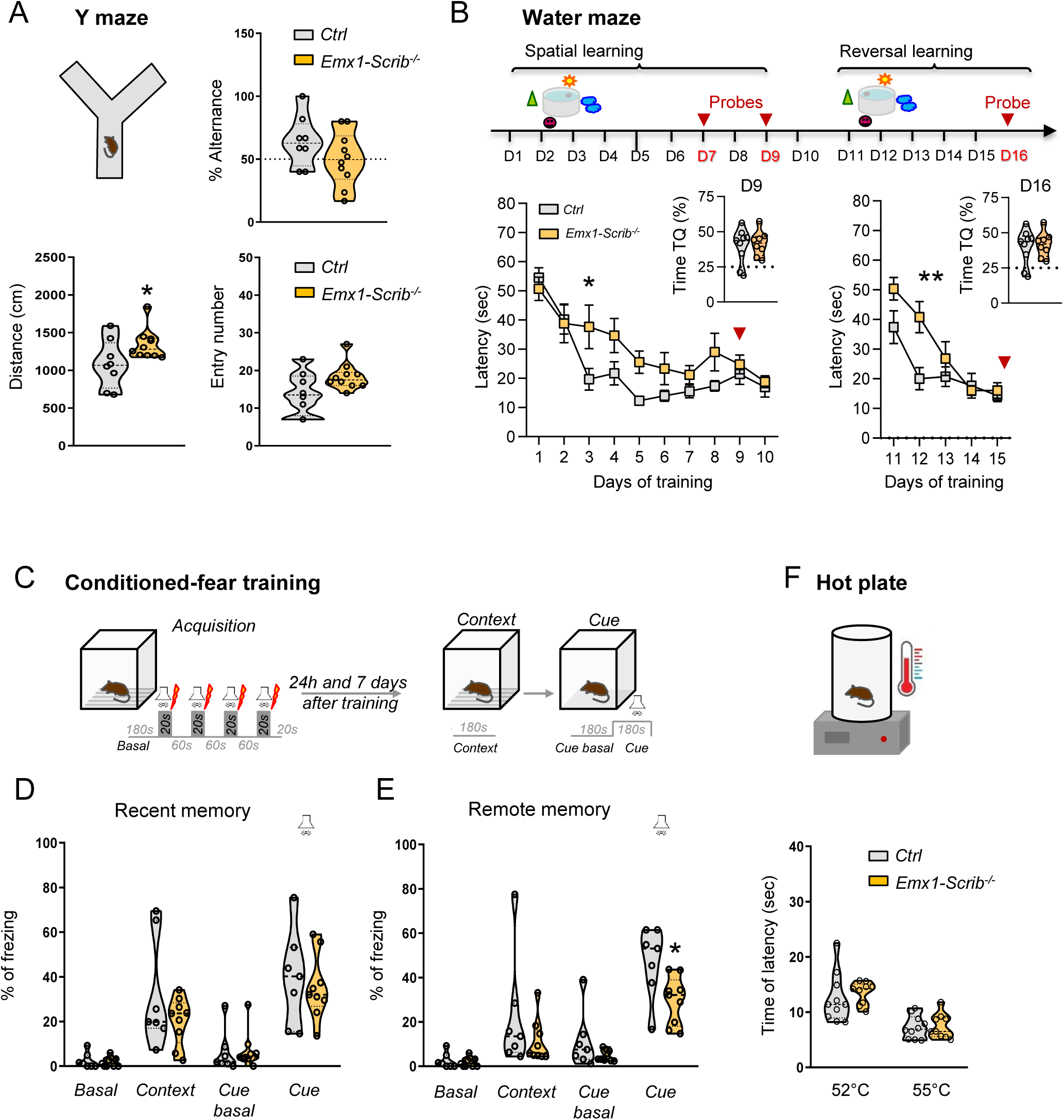
Spatial learning and long term memory deficit in Emx1-Scrib^-/-^ mice. **A**, Upper panel, Y maze test apparatus and alternance rate percentage in the Y maze test. Lower panel, total distance travelled and total number of entries in the Y maze test (control: n=8 mice; *Emx1-Scrib*^-/-^ : n=10 mice). **B**, Upper panel, Design of the Morris water maze apparatus task. Lower panel, the learning curve showing escape latency during the spatial and reversal training sessions. Time spent in target quadrant during the probe test at day 9 and day 16 are represented in the right corner of each graph (control: n=9 mice; *Emx1-Scrib*^-/-^ : n=8 mice). **C**, Upper panel, Design of the fear conditioning apparatus task. **D-E**, Freezing levels in context and tone fear conditioning. **D**, Percentage of Freezing measured before training (basal), 24 hours (recent memory) after training. **E**, Percentage of Freezing measured before training (basal) and 7 days (remote memory) after training (control: n=7 mice; *Emx1-Scrib*^-/-^ : n=9 mice). **F**, Lower, Hot plate apparatus. Time spent to paw withdrawal in the hot plate test at 52°C and 55°C (control: n=10 mice; *Emx1-Scrib*^-/-^ : n=9 mice). Data were represented as mean±SEM.

Considering that cKOs mice had impaired hippocampal and cortical connectivity, we examined whether the *Emx1-Scrib*^-/-^ mice were impaired in different types of learning and memory tests. In response to novelty measured over 2 hr, *Emx1-Scrib*^-/-^mutant mice showed a decrease in exploratory activity over time as the context loses its novelty, and ended up no different from control mice, suggesting that this simple form of spatial recognition is preserved in *Emx1-Scrib*^-/-^ cKO mice (time effect: *F*_3,532_=50,56, *P*<0.001***; **Fig 7C**). *Emx1-Scrib*^-/-^ mice showed the percentage of alternance comparable to that of WT littermates, suggesting that working memory assessed in Y maze is intact in the mutant mice (*t*-test: *t*_*16*_=1,404, *n*.*s*; **Fig 8A**). To further probe hippocampus-dependent connectivity, we next analyzed the effect of early *Scrib* loss on spatial learning and memory in the Morris water maze test. Mice were trained to learn spatial cues around the maze to find a hidden platform under the water during training sessions (training days: *F*_*9,135*_*=17*.*59, P<0,0001\*\*\**; **Fig 8B**). After training the mice for 15 days and preformed probe tests at day 7, 9 and 16, we observed that *Emx1-Scrib*^-/-^ mice showed significantly longer latency to find the platform during training sessions, demonstrating that the spatial learning is impaired in the mutants (genotype effect: *F*_*1*.*15*_*=5*.*53, P<0,05\**; **Fig 8B**). Furthermore, in the reversal test the *Emx1-Scrib*^-/-^ mutant mice showed a delay of spatial learning (interaction effect: *F*_*4*.*60*_*=4*.*335, P<0,01\*\**; Day2 *t*-test: *t*_*13*_=3.2, *P<0*.*05*;* **Fig 8B**). In all probes test, *Emx1-Scrib*^-/-^ mice show normal hippocampus-dependent memory. Importantly, although the *Emx1-Scrib*^-/-^mutant mice were hyperactive, swimming speeds during the probe tests were not different between genotypes. Finally, hippocampus-dependent context and amygdala-dependent tone associative memory was assessed by using classical fear conditioning paradigm in which the animals have to associate environmental cues to an electric shock (**Fig 8C**). Two different groups of *Emx1-Scrib*^-/-^ and control mice were re-exposed to the same environment 24 hours or 7 days after training, respectively showing the recent (24h) and the remote (7 days) contextual fear memory (**Fig 8D-E**). During the acquisition phase (without shock), all groups of mice displayed a normal freezing behaviour and the first 3 min of acquisition period was considered as the baseline period (basal). Every group exhibited an increase in freezing during the context and tone presentation tested 24 hours (test effect: *F*_*1*.*28*_*=75*.*69, P<0,0001\*\*\**; **Fig 8D**) and 7 days after training (test effect: *F*_*1*.*28*_*=75*.*82, P<0,0001\*\*\**; **Fig 8E**). The freezing levels were comparable between genotypes when the memory was tested 24 hours after training, showing that the recent contextual and tone fear memories are intact in *Emx1-Scrib*^-/-^ mice (no genotype effect: *F*_*1*.*56*_*=1*.*351, n*.*s*; **Fig 8D**). When mice were tested 7 days after training, *Emx1-Scrib*^-/-^ mice froze less than their control littermate during the tone indicating an acceleration of the extinction of the long term cued fear memory (genotype effect: *F*_*1*.*56*_*=8*.*459, P<0,01\*\**; **Fig 8E**). Here, *Emx1-Scrib*^-/-^ mutant mice exhibited no significant prolonged latency to paw to licking/jumping in the hot plate test than controls indicating normal nociceptive reactivity (no genotype effect: *F*_*134*_*=0*.*37, n*.*s*; **Fig 8F**).

Altogether, these results support the idea that *Emx1-Scrib*^-/-^ mutant mice display hyperactivity in novel and familiar environments, which is compatible with the altered psychomotor behavior observed in VRJS patients (see discussion). In addition, the remote memory deficits we observed in this mouse model have not been reported in patients with VRJS and should guide memory evaluation in older patients.

## Discussion

Our present study demonstrates that *Scrib* is essential for embryonic brain development and function. We showed that early deletion of *Scrib* leads to microcephaly and cortex layering defects associated with corpus callosum and hippocampal commissure agenesis. Behavioral analysis showed an increased locomotor activity accompanied by memory defects as a consequence. Our integrative work supports the participation of *Scrib* to various congenital neurodevelopmental deficits.

Both conditional mutants used in this study and with early deletion of *Scrib* (*FoxG1-Scrib*^-/-^ and *Emx1*-Scrib^-/-^ mutant mice) display microcephaly in a range that is comparable to well-established microcephalic mouse models ^35^. Microcephaly observed in mutants with embryonic *Scrib* deletion is most likely the result of a combination of disrupted neurogenesis, cell fate and migration processes, as deficits in any of these critical mechanisms can participate in brain malformation ^2^. Our data show that this phenotype arises from a depletion in the neuronal lineage, as observed by a decrease in CuxI- and Satb2-positive (layer II/III) CPNs, but also in Ctip2-positive (layer V) CPNs. The deletion of *Scrib* is accompanied by cortical layering defects with mispositioned neurons from layer V, suggesting migration defects that were validated by acute *Scrib* knockdown. This multifactor effect of *Scrib* is consistent with its expression profile. It is expressed in the cerebral cortex, both in glutamatergic neurons and in the radial glia, also supporting a role in migration. However, we also observed Scrib enrichment in the VZ and SVZ and we cannot rule out the possibility that it can contribute to asymmetric cell division (ACD) and cell fate determination, as observed in other systems ^6,7^.

The other main phenotype we observed after early *Scrib* deletion was the CC and DHC agenesis. These commissural defects may arise from cell fate, migration, and/or axonal outgrowth/guidance alterations. *Scrib* inactivation resulted in complete ACC in both rostral and caudal domains in *FoxG1-Scrib*^-/-^ cKOs, while *Emx1-Scrib*^-/-^ cKOs display ACC only in the caudal part. The most likely explanation for this restricted phenotype at the caudal level is that *Emx1* is expressed on a high-caudal to low-rostral gradient ^36^. In the rostral telencephalon, *Scrib*-dependent ACC is associated with the formation of abnormal swirls of axons called Probst bundles ^34^. Such phenotype is associated with failure of the callosal axons to cross the midline rather than an outgrowth problem towards the midline. Although we cannot completely exclude an autonomous role of *Scrib* in axon guidance as shown during axonal misrouting at the chiasm ^37^ and during spinal commissures formation in zebrafish ^38^, *Scrib* expression in the midline glia and the disorganization of that structure in *Emx1-Scrib*^-/-^ mutant mice suggest that a non-autonomous *Scrib*-dependent mechanism affects the crossing of the axons of the CC. Mutations in guidance cues expressed by the midline glia typically lead to a Probst bundles phenotype; thus, we can envision that *Scrib* deletion affects midline glial cells maturation and/or guidance cues secretion ^39^ as seen in other systems ^40,41^. Alternatively, by impacting these astroglial structures that are essential during interhemispheric remodeling ^42^, *Scrib* may promote tissue fusion ^43,44^, which in turn allow the callosal fibers crossing. Tissue fusion appears as a common theme during neural tube closure ^45^ and during interhemispheric remodeling in CC formation ^46^. Of note, one third to half of patients with spina bifida also have CC abnormalities ^27^ that may explain some behavioral deficits due to improper interhemispheric transfer of information ^28^.

By impairing cell fate, migration and commissure formation, we show that the absence of *Scrib* during development causes profound defects in the cortical layering and interhemispheric connectivity that underlie cognitive disabilities that are typical of neurodevelopmental disorders and some human syndromes. Edwards and collaborators discriminate two categories of human syndromes: on one hand the ones that display only a microcephalic phenotype and, on the other hand, the ones that do encompass both ACC and microcephaly ^3^. *SCRIB* may also fall into the latter category, reflecting a concomitant function for this gene in cortical organization and axonal guidance during development. Our results support the theory that the microcephaly and ACC observed in VRJS patients is the result, at least partially, of *SCRIB* haploinsufficiency.

Behavioral analysis of the *Scrib* cKO mice revealed alterations in psychomotor behavior. Specifically, *Emx1-Scrib*^-/-^ mutant mice had an increased locomotor behavior that is comparable to established hyperactive models ^47^. Because of the early and broad expression of *Emx1* in the brain, the pathological origin for the altered locomotor behavior may stem from the cortex and/or the hippocampus, through their connections. Exploration of cognitive performance shows that *Emx1-Scrib*^-/-^ display memory deficit for a cortex-dependent remote-cued fear memory recall. The CC primary function is to integrate motor, sensory, and cognitive activity between the two hemispheres ^3^. The DHC provides interhemispheric connections between hippocampi and despite few studies of the function of the DHC, the ability of the hippocampus to communicate effectively with contralateral homologous regions via the DHC may be important for cognitive performance, compounded by the other deficits observed in the mutants. This lack of communication and/or the microcephaly and/or layering defects could be the cause of these cognitive deficits. At the molecular level, it is tempting to speculate that Scrib may control some aspects of brain architecture and behavior, through known partner such as GIT1^48^ or Vangl2^49^. Similar to our model, mice lacking GIT1 have microcephaly ^50^ and display a hyperactivity phenotype combined with learning and memory deficits ^51^.Like *Emx1-Scrib*^-/-^ mice, *Emx1-Vangl2*^-/-^ display partial hippocampal and CC agenesis (but no microcephaly) that is caused by abnormal axonal outgrowth ^52^. However, a conditional deletion of *Vangl2* in postmitotic hippocampal granule cells does not alter spatial memory ^53^, highlighting the complexity of such molecular integration at the behavioral level. Additional work from our group support a critical role for Scrib in synaptic dysfunction and human psychiatric disorders ^13,54,55^, at least in part, by the regulation of the glutamatergic signaling ^18^. Remarkably, the function of this pathway in the process of learning and memory seems to extend toward the invertebrate phylum where *Scrib* is pivotal to the regulation of active forgetting ^56^. Future studies exploring these possibilities are needed to define the detailed molecular mechanisms.

Our findings uncover an essential role for *Scrib* in mammalian forebrain development and connectivity, both of which ultimately affect animal behavior. Given that several aspects of the neurological manifestations of VRJS were recapitulated in *Scrib* cKO mice, we suggest that *Scrib* may participate in this syndrome. The minimal common deletion found encompassed 3 genes including *SCRIB* and *PUF60* and displayed most of the cardinal features of VRJS ^29^. Although *PUF60* appears to be a major driver of VRJS syndrome ^29,57–64^, neurological features were reported in a much lower proportion, indicating that *PUF60* CNVs may not be their sole cause. From the human genetics standpoint, any VRJS-related phenotype due to *SCRIB* loss may prove difficult to observe because most *SCRIB* mutations lead to NTDs so deleterious that they may obscure more “subtle” phenotypes ^20–23^. The scarce number of NTD cases carrying *SCRIB* variants or VRJS patients implies the possibility that their true phenotype spectrum may be wider than indicated by publications providing few or no detailed neuropsychiatric evaluation. Patients with spina bifida tend to show altered cognitive abilities with language, memory, motor and psychosocial difficulties ^65^.

Neurological features for VRJS patients include mild intellectual disability, delay of developmental milestones such as standing upright and walking, delayed speech, feeding issues and generalized seizures ^66,67^. All of these features, combined with neuroanatomical features such as microcephaly and CC agenesis can fall under the umbrella of a neurodevelopmental disorders whose affected individuals develop psychomotor deficits reminiscent as those observed in ASDs, ADHD and schizophrenia. Interestingly, a case for a patient with ADHD revealed a chromosomal translocation breakpoint at 8q24.3 has been reported ^68^ and this region appears to overlap with ASDs as well ^32^. Further studies are warranted in order to determine whether other behavioral phenotypes such as seizure susceptibility or deficits of attention/impulsivity, are also recapitulated in *Scrib*^-/-^ cKOs. Our study demonstrates that *Scrib* mutant mice can provide an entry point to study its forebrain contribution in neuroanatomical and behavioral deficits observed in NTD and VRJS patients. Future investigation using both heterozygous *Scrib* KO and *Puf60* KO mice (alone or in combination) as a model is warranted to address their respective contribution and potential interaction in this syndrome at a more systemic level.

## Materials and methods

### Animal care and use

All mice were housed in the animal facility of the Neurocentre Magendie, in polypropylene cages under controlled conditions (at 23 ± 1 °C, with lights on from 7:00 A.M. to 7:00 P.M.). Food and water were available *ad libitum*. For timed pregnancy, the morning in detection of vaginal plug was designated as embryonic day E0.5. This study was performed according to the European Communities Council Directives (2010/63/EU) and has been approved by the Animal Care and Use Committee (Bordeaux) under the numbers 5012016-A and 5012015-A.

### Generation of brain-specific *Scrib* conditional knock-outs

Scrib spontaneous mouse mutant *Circletail* causes severe brain and neural tube damages that result in neonatal lethality ^8^, precluding the analysis of the role of *Scrib* during forebrain development ^17^. In order to circumvent this issue, we have applied a conditional gene-targeting strategy to inactivate *Scrib* at different developmental stages and in different cellular types in the brain. See Supplementary material and methods for details.

### Histology, *in situ* hybridization analysis and Immunofluorescence

For histology, heads (E16.5 embryos) or brains (P0 new-born) were harvested and fixed in Bouin’s or Carnoy’s fixative (Electron Microscopy Sciences) overnight as previously described ^69^. Samples were dehydrated in ethanol, paraffin-embedded, and sectioned (20 µm) coronally, horizontally or sagitally. Sections were collected onto Superfrost plus Gold slides (Thermo Scientific), stained with hematoxylin and mounted with Entellan (Millipore). The brains sections were examined using Leica MZ-16 stereomicroscope, imaged in the Bordeaux Imaging Center (http://www.bic.u-bordeaux.fr) using the NanoZoomer 2.0-HT slide scanner and analyzed using the Hamamatsu NDP viewer software (Hamamatsu). For *in-situ* hybridization, E16.5 and E17.5 embryos brains were dissected out, transferred in OCT solution and placed into the dry ice for storage at - 80°C before sectioning. *In situ* hybridization was performed using previously validated digoxigenin-labeled cRNA probes ^17^. 16 µm coronal embryonic brain sections of were postfixed in 4% paraformaldehyde / 0.2% glutaraldehyde for 10 min at room temperature (RT), bleached with 6% H2O2, digested in Proteinase K (5 µg/ml in PBS) for 2.5 min and postfixed in 4% paraformaldehyde / 0.2% glutaraldehyde for 10 min at RT. Then, slides were acetylated into freshly prepared 0.1M triethanolamine / PBS, pH 8, for 10 min at room temperature; 0.25% acetic anhydride acid was added for the last 5 min. Between each step, slides were rinsed with PBS. All subsequent steps were performed as previously described in ^17^. Images were acquired with Nanozoomer (Hamamatsu). For immunofluorescence staining (IF), P0 pups were anesthetized with pentobarbital, then perfused transcardially with 4% paraformaldehyde (PFA) in PB buffer (0.1% PBS, 0.9% NaCl, PH = 7.4). Dissected brains were postfixed in 4% PFA overnight at 4°C, infused in 30% sucrose in PB overnight and embedded in O.C.T (Sakura Finetek). The OCT-embedded brains were cryosectioned coronally at a thickness of 20 μm, mounted on Superfrost plus slides (VWR), washed with PBS, and immunostaining was performed. Sections were hydrated with PBS, permeabilized with 0.2 % Triton-X100/PBS (PBS-T), blocked using 10% Normal Goat Serum (NGS) and incubated overnight with the following primary antibodies in 5% NGS overnight at 4°C: rabbit anti-GFAP (DAKO; #Z0334;1:1000), rat anti-L1-CAM (Millipore; #MAB5272; 1:1000), mouse anti-RC2 (DSHB; 1:100), rat Ctip2 (Abcam; #ab18465, 1:500), mouse Satb2 (Abcam; #ab51502, 1:100), rabbit Tbr1 (Millipore; #AB9616; 1:500), rabbit Cux1 (Santa Cruz Biotechnology; #sc-13024; 1:100), rabbit anti-Scrib (homemade AbMM468; 1:300) ^70^. For Scrib immunostaining, brains were not perfused but only briefly fixed in 4% PFA for 30 min. Samples were washed three times in PBS-T and were incubated for 2 hr with the secondary antibodies Alexa Fluor® 488 or 546 Goat Anti-Mouse/Rat/Rabbit IgG (Life Technologies; 1:200) and then with DAPI (Life Technologies; 1:20000) for 30 min. Finally, samples were washed three times in PBS and mounted with Prolong Gold anti fading reagent (Invitrogen). Immunostained sections were imaged at similar brain region using a Zeiss Axio Imager Z1 microscope and Axio Vision (Version 4.7) imaging analysis software (Carl Zeiss). All images were processed with Photoshop CS5 software (Adobe) and ImageJ software (http://imagej.nih.gov).

### UDisco

Eight to ten-week old *Emx1-Scrib*^-/-^ mice received unilateral stereotaxic microinjections of a AAV (0.5/1µl at 300 nl per min) expressing GFP under the control of the promoter hSyn1 (AAV9-hSyn1-GFP-2A-GFP-f, titer 4.8 x10^12^ GC/ml) in the sensory-motor cortex region (anteroposterior [Y] -2 mm from Bregma, mediolateral [X] ± 1,2 mm, dorsoventral [Z] -0,5 mm) or expressing mCherry under the control of the CamKII promoter (AAV2/9-CamKII(0,4)-mcherry-WPRE, Vector Biolabs, titer 1.2 x1013 GC/ml) in the CA3 region of hippocampus (anteroposterior [Y] -2 mm from Bregma, mediolateral [X] ± 3 mm, dorsoventral [Z] -2,5 mm). Four weeks after surgery, the animals were perfused transcardially with PB followed by 4% PFA in PBS; the brains were removed and postfixed in 4% PFA for 24 hrs at 4°C and maintained in PBS. The entire brains were cleared using the uDISCO technique as described ^71^. The ultramicroscopy was performed using the system from LaVision BioTec (Bielefeld, Germany) equipped with a Fianium white laser, a sCMOS Andor camera, and a 0.5 NA 2X objective with a deeping lens. A zoom from 0.63 to 6.3 could be applied. Images and 3D reconstruction were analyzed with Image J and Imaris software.

### *In-utero* electroporation and tissue processing

*In utero* electroporation experiments were performed according to protocols previously described ^72^. The Animal Care and Use Committee (Bordeaux) has approved the experimental procedure under the number 5012015-A. Pregnant Swiss CD-1 mice were anesthetized using 4 % isoflurane in an anesthesia induction chamber, maintained with 2 % isoflurane with an anesthetic mask and injected before surgery with buprenorphine. Mice were subjected to abdominal incision; uterine horns were exposed and E14.5 embryos were placed on humidified gauze pads. Plasmid DNA was purified on Qiagen columns (EndoFree Plasmid Maxi Kit), resuspended in sterile endotoxin-free buffers (Qiagen) and mixed with Fast Green (Sigma). mCherry plasmid, together with pSuper or validated pSuper-Scrib shRNA construct (0.5 μg/μl) ^17^ were microinjected at a 1:3 ratio into the lateral ventricles of embryos. Five current pulses (50 ms pulse / 950 ms interval; 35–36 V) were delivered across the heads of the embryos using 7 mm electrodes (Tweezertrode 450165, Harvard Apparatus) connected to an electroporator (ECM830, BTX). Surgical procedure was completed with suture of the abdomen wall and skin. E18.5 embryos or P0 pups were processed for tissue analysis and immunostaining as described in the histology section. Subregions of the cerebral cortex (VZ/SVZ, IZ, LL and UL) were identified based on cell density using DAPI staining (Life Technologies; 1:20000). For each condition, sections from three embryos obtained from three separate litters were quantified. Quantification of mCherry-positive cells was performed using the cell counter plugin for ImageJ (http://rsbweb.nih.gov/ij/plugins/cell-counter.html). Data are given as a percentage of the total of cells positive for mCherry in each cortical subregion (mean ± SD).

### Behavioral testing

Behavioral experiments were conducted with *Emx1-Scrib*^-/-^ cKOs and their control littermate male mice of 10–11 weeks of age at the start of behavioral tests. All behavioral experiments were performed during the light phase (between 9:00 A.M. and 7.00 P.M.) of a 12 h light/dark cycle, under conditions of dim light and low noise. One week before starting the experiments, mice were housed in individual cage. Several cohorts of animals and multiple behavioral tests were used. Whenever possible, naïve animals were employed for behavioral testing; when the same cohort was used for multiple tests, the most stressful assays were performed last, to minimize between-test interference. To look for behavioral abnormalities mice were tested in activity cages (to measure locomotor activity); in elevated plus maze, open field and Y-maze (to measure exploratory activity and anxiety-like behavior); in rotarod, hot plate, beam walking and grid handling test (to measure sensory-motor activity); and in the fear conditioning test maze (to measure early and remote contextual end cued memory performance). All experimental apparatuses were cleaned with hydroalcoholic solution (Phagospray-DM) between subjects to remove odor residuals. See Supplementary material and methods for details.

### Quantification, statistical analysis and data representation

Details of statistical analyses and n values are provided in the figure legends subsections referring to individual assays. Statistical analyses were carried out using the GraphPad Prism statistical package (GraphPad). Normality of distribution and homogeneity of variance were validated and unpaired Student’s two-tailed t-tests for two data sets were used to compare groups with similar variance and are indicated along the P values in figures. P≤0.05 was considered as statistically significant. Statistics were derived from at least three independent experiments and not from technical replicates.

For behavior analyses, two-way ANOVA testing or repeated measure ANOVA was used for the evaluation of the effect of genotype and time in actimetry, motor activation, rotarod and fear conditioning behavioral test. The Bonferroni posthoc test was used when appropriate. The Student’s t-test was used for comparing genotype in other behavior tests. Whenever adequate, individual data points were reported as scatterplots to provide full information about the variability of data sets.

## Acknowledgements

We thank Neal Copeland, Nancy Jenkins (MD Anderson Cancer Center, USA) for providing *Scrib*^*fl/fl*^ floxed mice. We thank Anne Quiedeville, Chantal Médina and Ronan Peyroutou for their technical assistance. We also thank the "animal and genotyping facilities" members of the Neurocentre for technical assistance, notably Helene Doat, Sara Laumond, Claire Lordan and Delphine Gonzales and co-workers, but also the “Biochemistry and Biophysics Facility” of Bordeaux Neurocampus. We are grateful to Christel Poujol and Sébastien Marais, members of the Bordeaux Imaging Center (BIC, http://www.bic.u-bordeaux.fr), a service unit of the CNRS-INSERM and Bordeaux University, member of the national infrastructure France BioImaging supported by the French National Research Agency (ANR-10-INBS-04). These research facilities are funded by the Labex B.R.A.I.N. (ANR-10-LABX-0043). We thank Fatiha Boukhtouche and Peter Scheiffele for help with initial experiments using *in utero* electroporation. We thank all members of the “Planar Polarity and Plasticity” team, Fanny Mann and Valérie Castellani for thoughtful discussions.

## Author Contributions

Conceptualization: J. E., M. M. M., N. S., M. M.; Data Curation: J. E., M. M. M., T. M. M., M. S., M. D., M. R., F. C. de A.; Formal Analysis: J. E., M. M. M., T. M. M., M. S., M. D.; Funding Acquisition: J. E., N. S., M. M. ; Investigation: J. E., M. M. M., T. M. M., M. S., M. D., M. R., R. P., F. C. de A. ; Methodology: J. E., M. M. M., M. R., F. T., F. C. de A., N. S., M. M. ; Project Administration: J. E., F. T., F. C. de A., N. S., M. M. ; Resources: J. E., M. M. M., M. R., R. P., R. R., F. T., F. C. de A., N. S., M. M. ; Supervision: J. E., F. T., F. C. de A., N. S., M. M. ; Validation: J. E., M. M. M., T. M. M., M. S., M. D., M. R., F. C. de A., N. S., M. M. ; Visualization: J. E., M. M. M.; Writing – Original Draft Preparation: J. E., M. M. M., N. S., M. M. ; Writing – Review & Editing: J. E., M. M. M., T. M. M., M. S., M. D., F. T., F. C. de A., N. S. and M. M.

## Funding

This research was supported by INSERM, the French National Research Agency ANR NeuroScrib ANR-07-NEUR-031-01 (NS, MM), La Fondation pour la Recherche Médicale FRM grant #SPF20101221210 (JE) and " Equipe FRM " grant #DEQ20160334899 (MM), the European Commission Marie Curie Action CIG #303820 (JE), but also ERANET-NEURON (BMBF 01EW1910), JPND (BMBF 01ED1806), and Deutsche Forschungsgemeinschaft (DFG grant CA1495/4-1) (FCDA).The Montcouquiol/Sans lab is member of the Labex B.R.A.I.N. The funders had no role in study design, data collection and analysis, decision to publish, or preparation of the manuscript.

**Correspondence and requests for materials** should be addressed to and/or

## Supplementary materials and methods

### Generation of brain-specific *Scrib* conditional knock-outs

*FoxG1*-Cre, *Emx1*-Cre and the Cre reporter mouse strain B6.Cg-*Gt(Rosa)26Sor*^*tm6(CAG- ZsGreen1)Hze*^/J (Ai6) mice ^1^ were purchased from The Jackson Laboratory (Bar Harbor, ME). *Scrib*^*fl/fl*^ floxed mice were generated previously in collaboration with Neal Copeland, Nancy Jenkins and Rivka Rachel at NIC/NIH ^2^. *Scrib* mouse gene contains 38 exons that are translated into a full-length 180 kDa protein. In *Scrib*^*fl/fl*^ floxed mice, lox-P sequences are inserted before exon 2 and after exon 8 ^3^. When crossed with a mouse line expressing the Cre recombinase, all the exons between 2 and 8 are excised, leading to the loss of the full length protein. Heterozygous rodents were intercrossed to generate homozygous and wildtype littermates as previously published ^4^. *Scrib*^*fl/fl*^ floxed mice were either crossed with *FoxG1*-Cre or *Emx1*-Cre mice to obtain compound mice, respectively referred to as *FoxG1-Scrib*^-/-^ and *Emx1-Scrib*^-/-^ mutant mice with early excision in different cellular types in the brain. All Scrib^-/-^ cKO mice are viable, fertile and do not display any gross physical abnormalities. Cre-mediated recombination in *FoxG1*-Cre mice starts as early as E8.5 and occurs mostly in all cells from both dorsal and ventral telencephalon ^5^. This includes progenitors from neuroepithelium that will specify various neuronal and glial cells from the neocortex and hippocampus, but also from the ganglionic eminence that will give rise to GABAergic interneurons. In *Emx1*-cre mice, the Cre recombinase is expressed later as compared with *FoxG1*-Cre mice, starting as early as E10.5 and only in the dorsal telencephalon ^6^. Emx1 pattern of expression lead to the excision of *Scrib* in the vast majority of the neurons of the neocortex and hippocampus, in glial cells of the pallium, but not in the GABAergic interneurons. Genotyping was performed as described previously by PCR using the following primers: F (5’-gcacactgggtatcatggcta-3’), R1 (5’-gcaatctccagagccttacaga-3’), R2 (5’-cccttggaaacctacatcccaa-3’) ^3^. Wild-type (WT), floxed, deleted *Scrib* alleles were distinguished by the following amplified products: for WT band (F+R1; 437 bp), flox band (F+R1; 541 bp) and cKO band (F+R2; 193 bp) **(Fig 2A)**. Cre genotyping was performed using the following primers: F (5’-cggcatggtgcaagttgaata-3’), R (5’-gcgatcgctattttccatgag-3’), resulting in a 300 bp band. Cre minus littermates showed no detectable phenotype and were used as controls. The body weight of Emx1-Scrib^-/-^ was slightly lighter than their littermate controls **(Fig S6A)**. It is worth noting that none of the *Scrib*^-/-^ mutants were homozygous for the knock-in *Emx1*-Cre allele (i.e., all were obligate Cre heterozygotes), thus avoiding potential confounds related to *Emx1* loss-of-function. The Cre reporter mouse strain B6.Cg-*Gt(Rosa)26Sor*^*tm6(CAG-ZsGreen1)Hze*^/J (Ai6) ^1^ which expresses ZsGreen1 from the *Rosa26* locus, was used for evaluation of Emx1-Cre-induced recombination efficiency. *Emx1-Scrib*^-/-^ mice were crossed with Ai6 reporter mice, resulting in enhanced green fluorescent protein (ZsGreen1) expression following Cre-mediated recombination selectively in Emx1-expressing cells within the cortex, hippocampus, fimbria and fibers within the caudate putamen **(Fig 2G-H)**. In contrast, the striatum (ZsGreen1-negative) lacked Cre activity, consistent with Emx1 expression pattern in the dorsal telencephalon.

### Quantitative analysis of laminar position in the cerebral cortex

Sections chosen for analysis were matched along the rostral-caudal axis and observed at the caudal level for every sample (where ACC is observed in Emx1-Scrib^-/-^ cKO brains). Immunofluorescence images were converted to gray values and normalized to background staining. Regions of interest (ROIs) with a fixed width (but of variable length, corresponding to the thickness of the cortex) were positioned in the motor cortex region with the long axis perpendicular to the pial surface. Each ROI was subdivided into ten equal bins from the pia (bin 1) to the outer border of the intermediate zone (bin 10) to assess CPN distribution across the layers of the cortex. In such arrangement, layers II-III (Cux1- and Satb2-positive) were corresponding to bins 2-4 while layer V (Ctip2-positive) was corresponding to bins 5-7 (highlighted in red in every figure). Slides were evaluated blindly to the genotype and the number and distribution of CuxI-, Satb2-, Ctip2- and DAPI-labeled cells in each zone was determined manually using the cell counter plugin for ImageJ (http://rsbweb.nih.gov/ij/plugins/cell-counter.html). In order to take into account a possible effect of Scrib deletion on total cell number in the cortex, data are given as a ratio (in percent) of the total of cells positive for each marker to DAPI-positive cells in each bin (mean ± SEM). Analysis was performed using at least 3 independent experiments (3 to 4 brains per genotype were quantified). The effect of the genotype on the distribution of cells within the bins was assessed using the Student test (*t test*).

### Callosal Axon tracing

Corticocortical tract tracing was performed as previously described ^7^ using fluorescent carbocyanine dye (1, 1′-Dioctadecyl-3, 3, 3′, 3′-Tetramethylindocarbocyanine Perchlorate; DiI, Invitrogen). Brains were dissected at P0 and fixed in 4% PFA. A single DiI crystal was inserted in the dorsomedial cortex and allowed to diffuse for 1 month. Coronal vibratome sections of 150 μm thickness were cut on a vibratome, mounted on slides and imaged as described in the previous paragraph.

### Western blot

Cerebral cortices of P0 cKO pups and their control littermates were homogenized in RIPA buffer (10 mM Tris-Cl (pH 8.0), 1 mM EDTA, 0.5 mM EGTA, 1% Triton X-100, 0.1% sodium deoxycholate, 0.1% SDS, 140 mM NaCl) supplemented with protease inhibitor cocktail (Complete™; Roche). Protein concentration was determined using Pierce BCA Protein assay kit (Thermo Scientific). Equal amounts of protein were diluted in sample buffer, separated by sodium dodecyl sulfate–polyacrylamide gel electrophoresis (SDS-PAGE), and visualized using the enhanced chemiluminescence (ECL) as previously described ^4^. Briefly, protein extracts were separated on an 8% SDS-PAGE and transferred overnight to polyvinylidene difluoride membranes (Millipore). Membranes were blocked with 5% nonfat milk in 1×Tris-buffered saline pH 7.4; 0.05% Tween-20 (TBS-T) for 30 min at room temperature (RT) and were probed with homemade rabbit anti-Scrib (MM468; 1:500) ^8^ and mouse anti-GAPDH (Millipore; 1:1000) for 1hr RT. Membranes were incubated with secondary antibodies (GE Healthcare UK; donkey anti-rabbit or anti-mouse IgG conjugated to horseradish peroxidase, 1:5000) in 1% nonfat milk in TBS-T for 1hr RT. Each incubation step was followed by 3 washes with TBS-T for 10 min and immunoreactive signals were detected using Pierce ECL substrate (Thermo Scientific). Band intensity was analyzed by densitometry with the ImageJ software (http://imagej.nih.gov) and quantified as a percentage of control band intensity using a representative of at least three independent experiments.

### *In-utero* electroporation and tissue processing

*In utero* electroporation experiments were performed according to protocols previously described ^9^. The Animal Care and Use Committee (Bordeaux) has approved the experimental procedure under the number 5012015-A. Pregnant Swiss CD-1 mice were anesthetized using 4 % isoflurane in an anesthesia induction chamber, maintained with 2 % isoflurane with an anesthetic mask and injected before surgery with buprenorphine. Mice were subjected to abdominal incision; uterine horns were exposed and E14.5 embryos were placed on humidified gauze pads. Plasmid DNA was purified on Qiagen columns (EndoFree Plasmid Maxi Kit), resuspended in sterile endotoxin-free buffers (Qiagen) and mixed with Fast Green (Sigma). mCherry plasmid, together with pSuper or validated pSuper-Scrib shRNA construct (0.5 μg/μl) ^10^ were microinjected at a 1:3 ratio into the lateral ventricles of embryos. Five current pulses (50 ms pulse / 950 ms interval; 35–36 V) were delivered across the heads of the embryos using 7 mm electrodes (Tweezertrode 450165, Harvard Apparatus) connected to an electroporator (ECM830, BTX). Surgical procedure was completed with suture of the abdomen wall and skin. E18.5 embryos or P0 pups were processed for tissue analysis and immunostaining as described in the histology section. Subregions of the cerebral cortex (VZ/SVZ, IZ, LL and UL) were identified based on cell density using DAPI staining (Life Technologies; 1:20000). For each condition, sections from three embryos obtained from three separate litters were quantified. Quantification of mCherry-positive cells was performed using the cell counter plugin for ImageJ (http://rsbweb.nih.gov/ij/plugins/cell-counter.html). Data are given as a percentage of the total of cells positive for mCherry in each cortical subregion (mean ± SD).

### Behavioral testing

#### Plus maze, Open field, Y maze and locomotor activity

Elevated plus maze, Open field and Y-maze experiments were performed as described previously ^3,10^. Locomotor activity in response to novelty and daily rhythm of activity experiments were performed as described previously ^10^.

#### Rotarod

Mice performance on the rotarod, which accelerates from 4 to 40 rpm in 5 minutes, was evaluated for 5 trials per session on three consecutive days. A resting time of 15 minutes was allowed between each trial. The end of a trial was considered when mice were falling off the rod. Latency to fall was recorded for each trial. The average latency was calculated for each testing day.

#### Hot plate

Animals were placed individually on a hot plate with the temperature adjusted to 49°C.Response latency (sec) to jump or lick the hind paws was measured. The cut off time was taken as 30 seconds to avoid risk of thermal injury to the skin.

#### Beam walking

During three successive days, mice were placed on one end of a wood beam (180 cm in length with a 2 or 1 cm square cross-section) at a height of 50 cm above a container with soft bedding and the time required to reach an enclosed safety box at the other end (80 cm away) is measured. On training day 1, beam was place horizontally and each mouse was trained to traverse the 2cm beam and then the 1cm beam. On training day 2, each mouse was trained to traverse the horizontal 1cm beam and then the 1cm beam inclined at an angle of 10° from ground. Tree training sessions par beam and the mice rest for 10 min in their home cages between training sessions on the two beams. On the day test, time taken to traverse the 1cm beam inclined at an angle of 10° from ground and the number of paw faults or slips are recorded. Analysis of each measure was based on the mean scores of the two best trials.

#### Grid handling

Mice were placed on a metal grid (spacing 1 cm2) and allowed to grip the grid with four paws. The grid was inverted at an angle of 180°, 20 cm above the ground and the time for the mouse to fall onto soft bedding was measured in seconds.

#### Morris water maze

Spatial learning and memory were performed as described previously ^10^ except for the reversal acquisition training where the platform was moved to the opposite quadrant used previously for the spatial learning test.

#### Fear conditioning

On day1, mice were trained in a standard fear conditioning apparatus. The training consisted of a single trial and was performed in a constantly illuminated Plexiglas cage (transparent walls and metal grid floor). Fear conditioning was performed by placing the mice in context A for 180s acclimation period followed by three conditional-unconditional stimulus (CS-US) separated by 120s interval. The CS-US consisted of 3 successive tones (CS, 20 s, 65 dB) and a footshock (US, 0.4 mA, 1 s, constant current) delivered through a stainless-steel grid floor. The fear conditioning chamber was thoroughly cleaned with 70% ethanol before each mouse was placed in the box. Memory for the context and the tone were evaluated on day 2 (recent memory) or 8 (remote memory) following conditioning. For the contextual memory test, mice were placed into the conditioning chamber and allowed to explore for 360 sec. For the tone memory test, the same mice were placed in a novel cage (cage with colored walls and flat plastic floor) 3 hr after the contextual memory test, allowed to acclimate to the chamber for 180sec and then presented with tone (180 sec, 65 dB). Freezing, an index of fear defined as the lack of movement except for heart-beat and respiration, was recorded during the 180sec acclimation period and context or tone presentation ^11^.

## Supplementary Figure legends

**Supplementary Figure S1.**
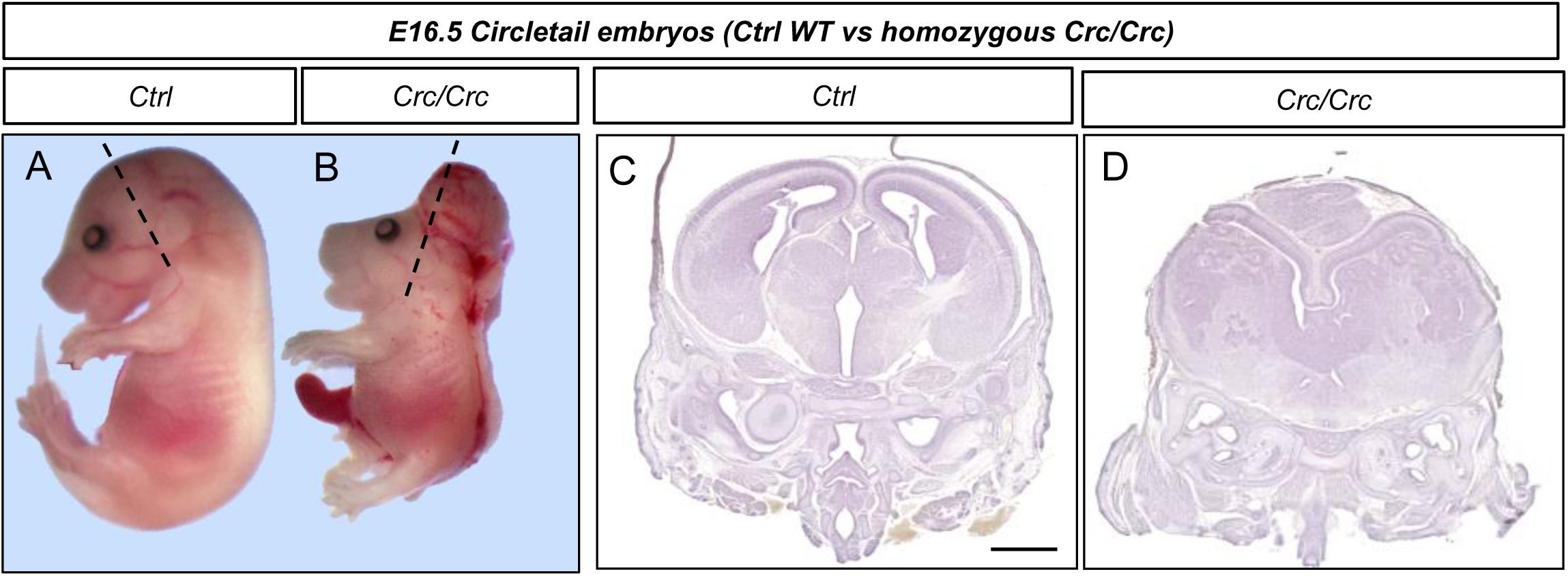
Circletail mutants exhibit severe neural tube defects / craniorachischisis, impeding brain development analysis. **A**, E16.5 control littermate and **B**, Circletail Crc/Crc heterozygous embryos in lateral view. Coronal sections of **C**, control and **D**, Crc/Crc mutant, performed at the level of the dashed lines in A, B, and stained with hematoxylin. Scale Bar: 100µm.

**Supplementary Fig S2.**
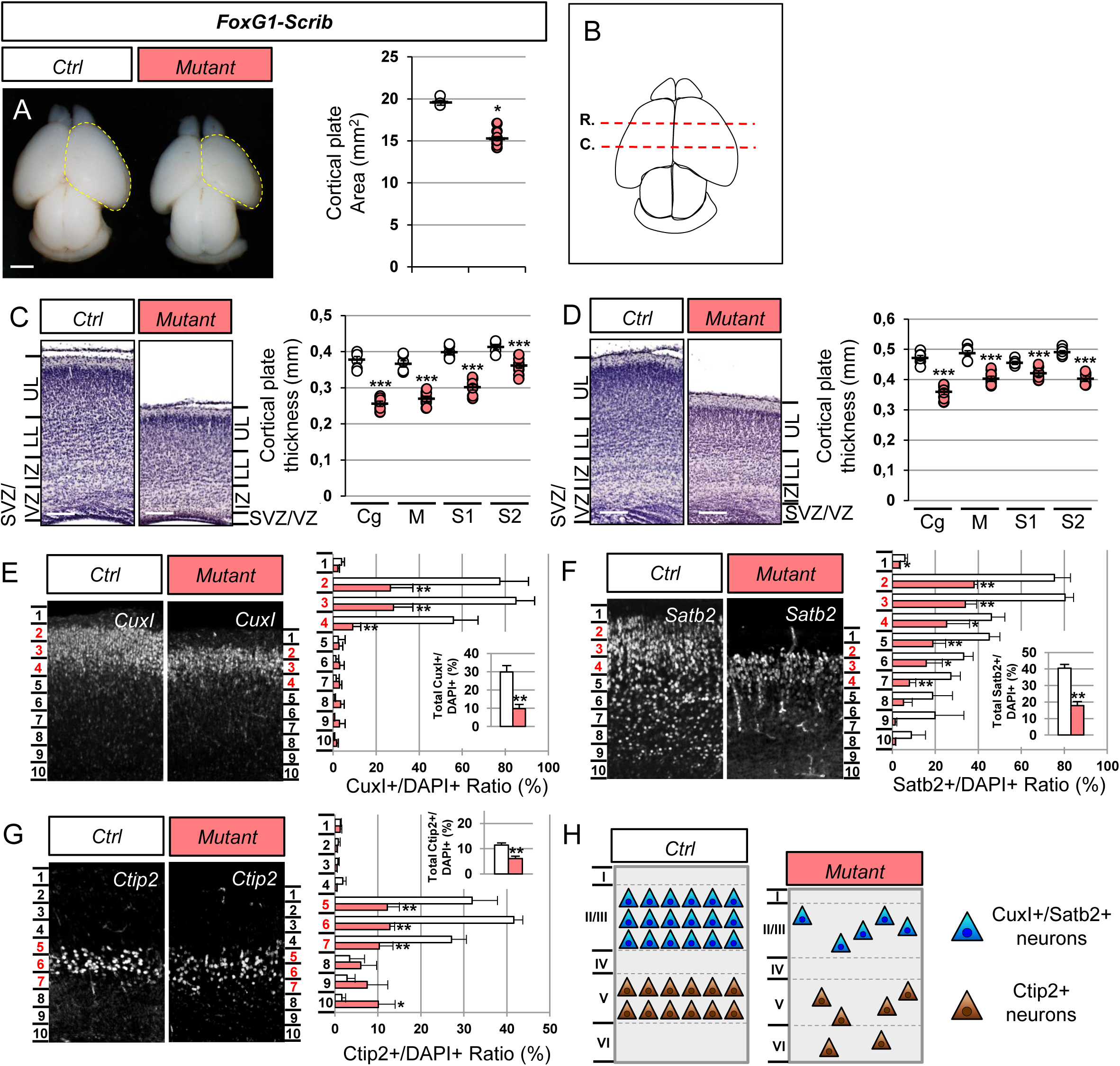
Microcephaly and severe cortical layering defects in *FoxG1-Scrib*^-/-^ cKO brains. **A**, Dorsal views of P0 *FoxG1-Scrib*^-/-^ cKO brains. Cortical plate areas are marked with a yellow dashed line. Statistical analysis via a two-tailed t test (*P*<0*.*05*) using between 5 and 8 brains per genotype from at least 3 independent experiments. Error bars indicate the SD. Scale bar: 1 mm. **B**, Schematic view of a P0 brain sectioned coronally at the rostral (R.) or caudal (C.) level. **C-D**, Representative hematoxylin staining of coronal sections from newborn *FoxG1-Scrib*^-/-^ cKO motor cortex at the caudal (C) and rostral (D) levels and their respective controls. A marked reduction of the motor cortex thickness (M) in cKOs extends to the cingulate (Cg) and somatosensory (S1-S2) cortex at both rostral (C) and caudal (D) levels. Cortical plate thickness was measured radially from the top of the upper layer (UL) to the bottom of the lower layer (LL) of the cortex. IZ: Intermediate Zone, SVZ: Sub-Ventricular Zone, VZ: Ventricular Zone. Statistical analysis via a two-tailed t test (*P***<0*.*0001*) using between 6 to 8 measurements per genotype from at least 3 independent experiments. Error bars indicate the SEM. Scale bar: 0.2 mm. **E-G**, Representative Immunofluorescence staining of CuxI (E), Satb2 (F) and Ctip2 (G) on coronal sections from newborn *FoxG1-Scrib*^-/-^ cKO brains in caudal motor cortex. Quantification of CuxI-, Satb2- and Ctip2-positive neurons is shown as a percentage (see methods). Severe reduction of CuxI (ctrl, 29.9% ± 3.4; mutant, 9.75% ± 2.3; *p* = 0.003), Satb2 (ctrl, 40.5% ± 2.2; mutant, 17.9% ± 2.4; *p* = 0.001) and Ctip2 (ctrl, 11.4% ± 0.8; mutant, 6.1% ± 0.9; *p* = 0.006) percentages were observed in *FoxG1-Scrib*^-/-^ cKO caudal motor cortices. Statistical analysis via a two-tailed t test (*P*<0*.*05, P**<0*.*01)* using between 3 to 4 measurements per genotype from at least 3 independent experiments. Error bars indicate the SD. **H**, Schematic representation of cortical layering in caudal motor cortex of *FoxG1-Scrib*^-/-^ cKO and its control. Both early-born (brown) and late-born neurons (blue) are massively decreased in proportion suggesting cell fate defects. See also Fig3.

**Supplementary Fig S3.**
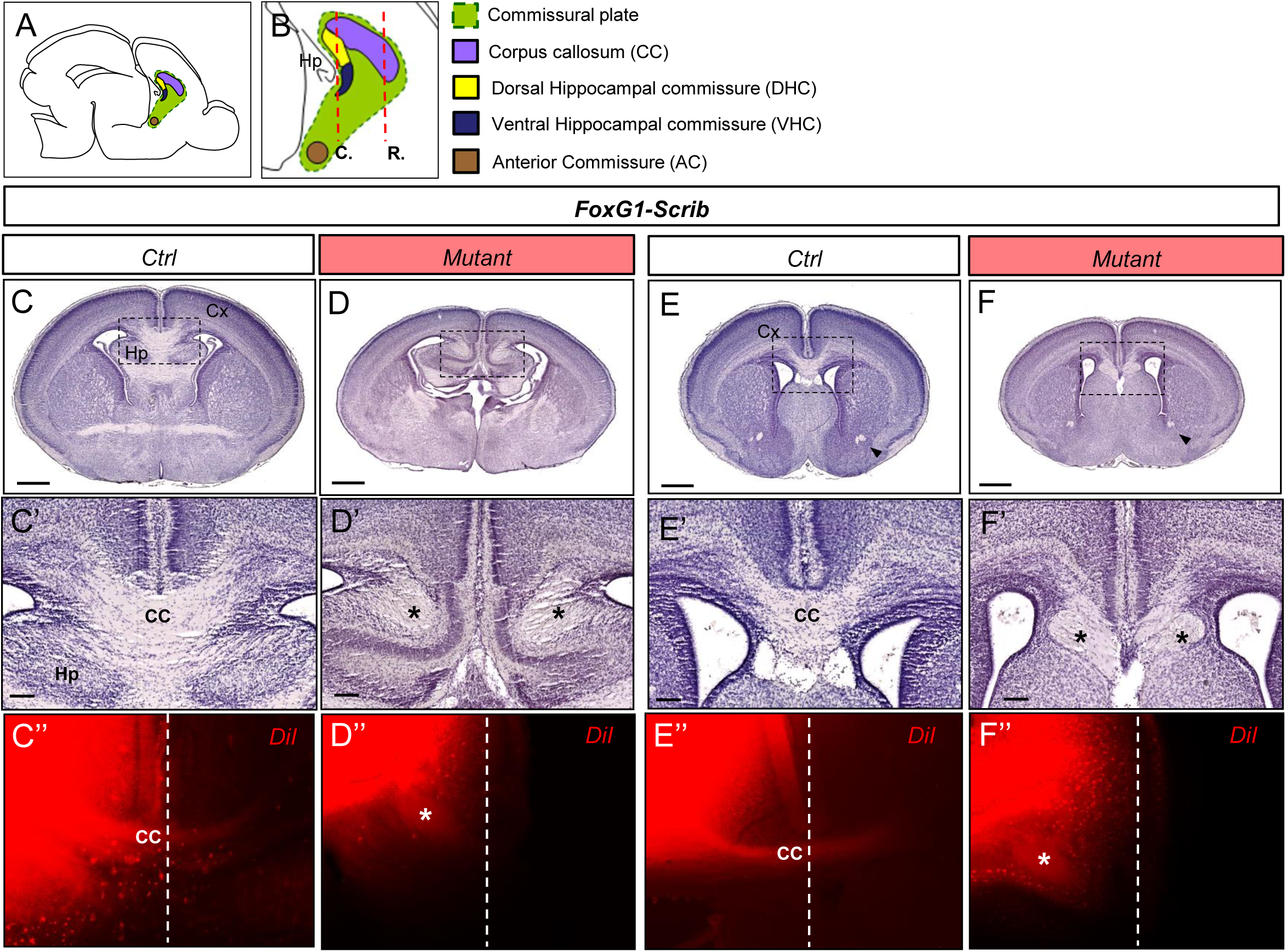
Complete corpus callosum agenesis in *FoxG1-Scrib*^-/-^ *cKO* mutants. **A-D**, Representative hematoxylin staining of coronal sections from newborn *FoxG1-Scrib*^-/-^ cKO brains (B and D) and their respective controls (A and C) at the caudal (A-B) or rostral (C-D) levels. **A’-D’**, Higher magnification for selected insets (boxed areas) from (A-D) illustrating high penetrance of CC agenesis (ACC) both at the caudal and rostral level. At P0, 100% of *FoxG1-Scrib*^-/-^ (n=12) cKO brains displayed ACC. Instead of crossing the midline, CC axons formed whorls (Probst bundles, PB) on either side of the midline that are indicated with an asterisks in B’ and D’. **A’’-D’’**, DiI crystals placed in the dorsomedial cortex trace CC axons in *FoxG1-Scrib*^-/-^ cKO brains (B” and D”) and their respective controls (A” and C”) at the caudal (A”-B”) or rostral (C”-D”) levels at P0. In *FoxG1-Scrib*^-/-^ cKO brains, misrouted callosal axons form dense PB (asterisks in B”, D”) lateral to the midline. Abbreviations: Cortex (Cx), Hippocampus (Hp), Corpus Callosum (CC). Midline is indicated as a white dashed line. Scale bars: 1 mm in (A-D), 0.1 mm in (A’-D’).

**Supplementary Fig S4.**
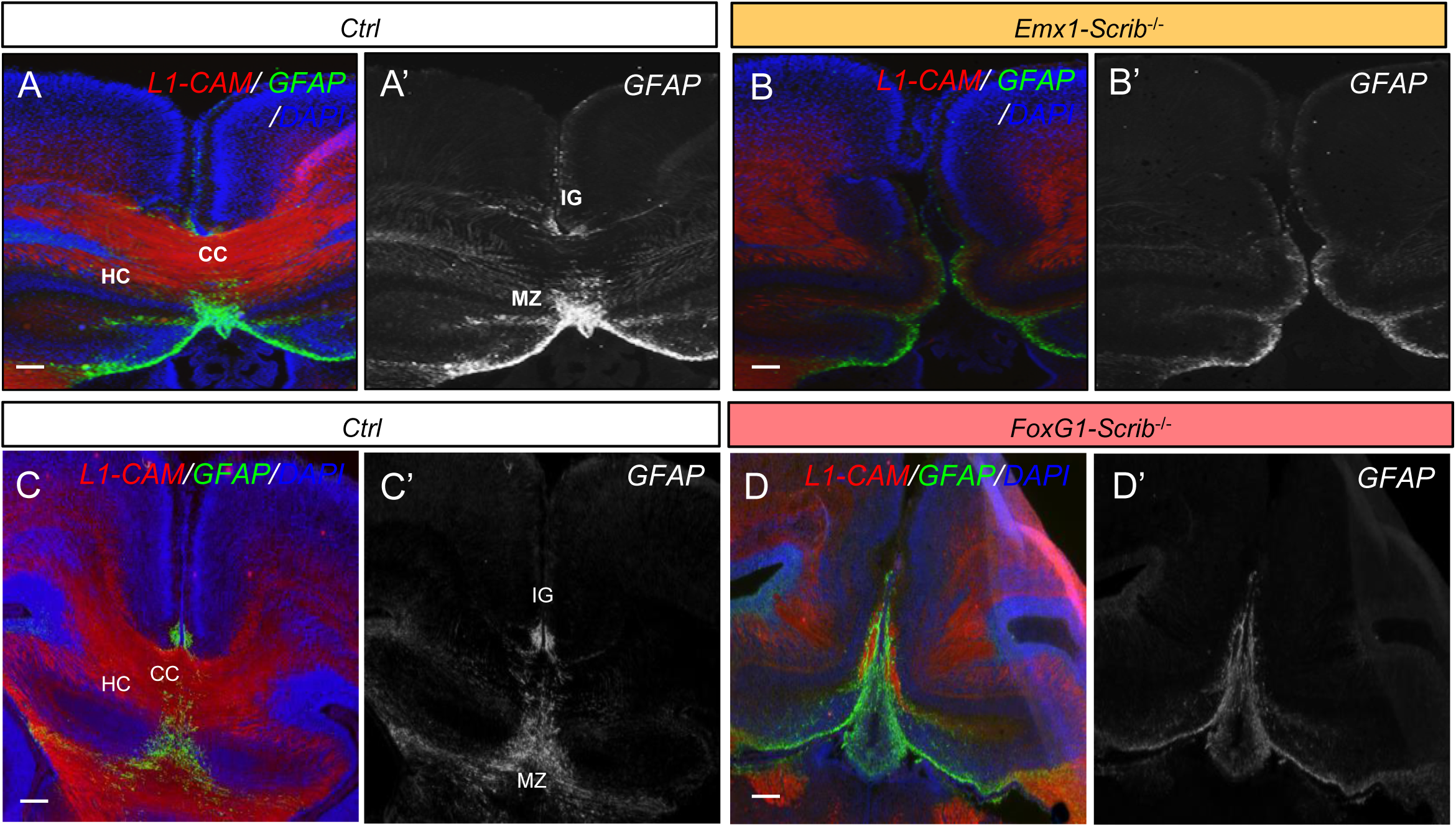
Corpus callosum agenesis in *Scrib cKO* mutants is associated with impaired midline glia structure positioning. **A-D’**, Representative immunofluorescence staining of L1-CAM (red) and GFAP (green) as a merged image (A-D) or GFAP only (A’-D’, gray) on coronal sections from newborn *Emx1-Scrib*^-/-^ (B and B’, orange) and *FoxG1-Scrib*^-/-^ (D and D’, red) cKO brains together with their respective controls at the caudal level. In *Scrib* cKO brains, agenesis of the corpus callosum is confirmed by the failure of L1-CAM-positive axonal fibers to cross the midline. GFAP-positive midline glial structures such as the MZ are missing in these mutants, resulting in improper hemisphere fusion and failure for the IG to form and localize appropriately. Scale bars: 0.1 mm in (A-D).

**Supplementary Fig S5.**
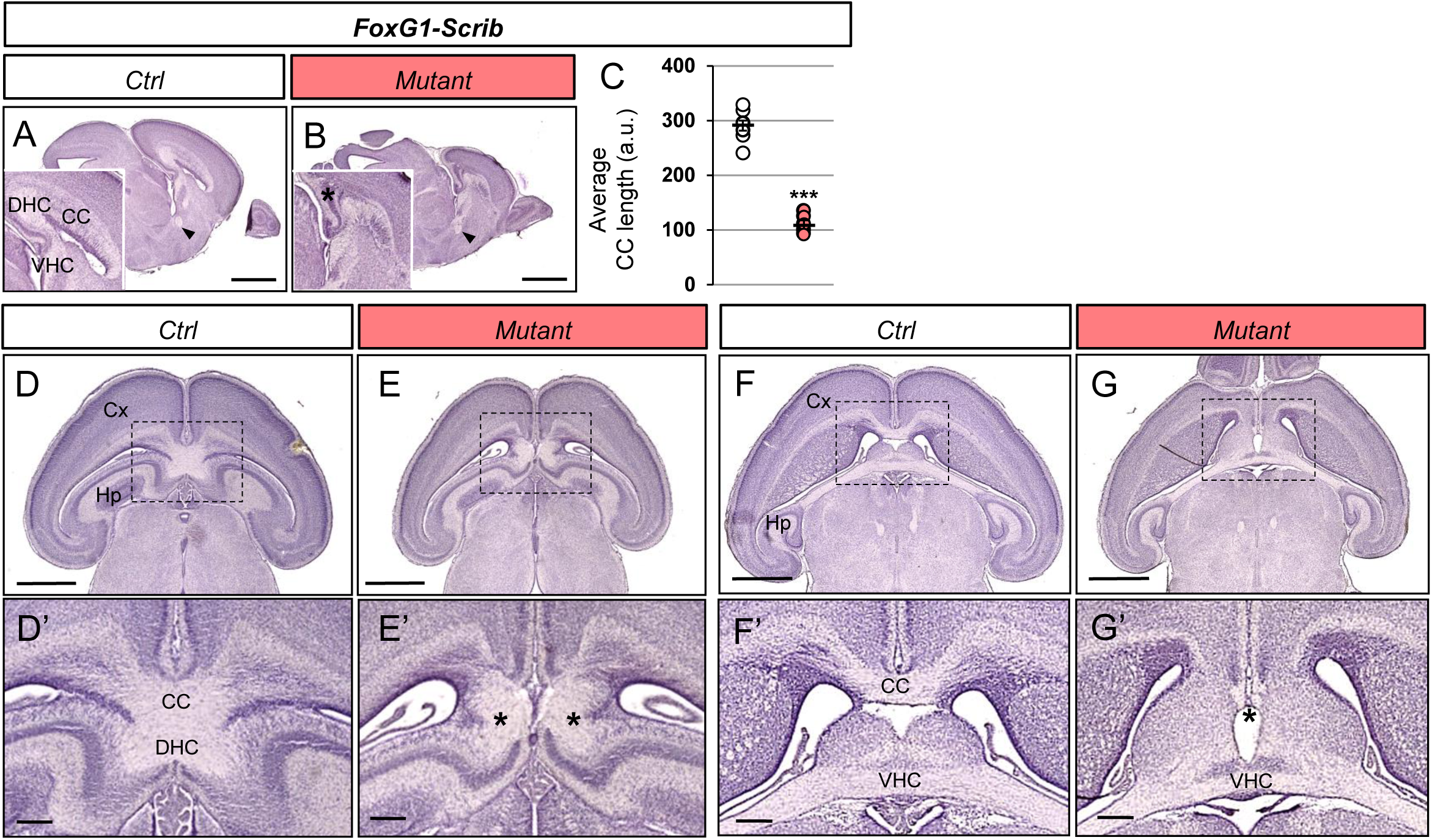
ACC is accompanied by hippocampal commissure agenesis in *Scrib*^-/-^ cKO mutants. **A-B**, Representative hematoxylin staining of para-sagittal sections from newborn *FoxG1-Scrib*^-/-^ KO brains (B) and its control (A). Commissural plates are magnified in each inset. The AC (indicated by arrowheads) is still present in brains from cKO mutants. **C**, Marked reduction of CC length (arbitrary units a.u) along the rostro-caudal axis in *FoxG1-Scrib*^-/-^ cKO brains (red, n=8) as compared with their control littermates (white, n=6). Statistical analysis via a two-tailed t test (*P***<0*.*0001*) using between 5 and 8 brains per genotype from at least 3 independent experiments. **D-G**, Representative hematoxylin staining of serial horizontal sections from newborn *FoxG1-Scrib*^-/-^ (E,G) and their respective controls (D,F) at the dorsal (D-E) and ventral level (F-G). **D’-G’**, Higher magnification for selected insets (boxed areas) from (D-G). Sections through the *FoxG1-Scrib*^-/-^ cKO brains reveal the complete absence of the CC and DHC. Misrouted callosal axons form dense PB (asterisks) lateral to the midline in dorsal sections and failed to reach the contolateral hemisphere ventrally (E’, asterisk). Despite an apparent thinning, the VHC is still observed in ventral sections (G’).These defects were penetrant in all *Scrib*^-/-^ cKO mutants observed (n=3 for each genotype). Scale bars: 1 mm in (A-B, D-G), 0.2 mm in (D’-G’).

## References

1. Lancaster, M. A. & Knoblich, J. A. Spindle orientation in mammalian cerebral cortical development. Curr. Opin. Neurobiol. 22, 737–746 (2012).

2. Evsyukova, I., Plestant, C. & Anton, E. S. Integrative mechanisms of oriented neuronal migration in the developing brain. Annu. Rev. Cell Dev. Biol. 29, 299–353 (2013).

3. Edwards, T. J., Sherr, E. H., Barkovich, A. J. & Richards, L. J. Clinical, genetic and imaging findings identify new causes for corpus callosum development syndromes. Brain 137, 1579–1613 (2014).

4. Iyer, J. & Girirajan, S. Gene discovery and functional assessment of rare copy-number variants in neurodevelopmental disorders. Brief Funct Genomics 14, 315–328 (2015).

5. Hu, W. F., Chahrour, M. H. & Walsh, C. A. The diverse genetic landscape of neurodevelopmental disorders. Annu Rev Genomics Hum Genet 15, 195–213 (2014).

6. Stephens, R. et al. The Scribble Cell Polarity Module in the Regulation of Cell Signaling in Tissue Development and Tumorigenesis. J. Mol. Biol. (2018) doi: 10.1016/j.jmb.2018.01.011.

7. Bonello, T. T. & Peifer, M. Scribble: A master scaffold in polarity, adhesion, synaptogenesis, and proliferation. J. Cell Biol. 218, 742–756 (2019).

8. Murdoch, J. N. et al. Disruption of scribble (Scrb1) causes severe neural tube defects in the circletail mouse. Hum. Mol. Genet. 12, 87–98 (2003).

9. Montcouquiol, M. et al. Identification of Vangl2 and Scrb1 as planar polarity genes in mammals. Nature 423, 173–177 (2003).

10. Ezan, J. & Montcouquiol, M. Revisiting planar cell polarity in the inner ear. Semin. Cell Dev. Biol. 24, 499–506 (2013).

11. Goodrich, L. V. The plane facts of PCP in the CNS. Neuron 60, 9–16 (2008).

12. Tissir, F. & Goffinet, A. M. Shaping the nervous system: role of the core planar cell polarity genes. Nat. Rev. Neurosci. 14, 525–535 (2013).

13. Sans, N., Ezan, J., Moreau, M. M. & Montcouquiol, M. Planar Cell Polarity Gene Mutations in Autism Spectrum Disorder, Intellectual Disabilities, and Related Deletion/Duplication Syndromes. in Neuronal and Synaptic Dysfunction in Autism Spectrum Disorder and Intellectual Disability 189–219 (Elsevier, 2016). doi: 10.1016/B978-0-12-800109-7.00013-3.

14. Wada, H. et al. Dual roles of zygotic and maternal Scribble1 in neural migration and convergent extension movements in zebrafish embryos. Development 132, 2273–2285 (2005).

15. Walsh, G. S., Grant, P. K., Morgan, J. A. & Moens, C. B. Planar polarity pathway and Nance-Horan syndrome-like 1b have essential cell-autonomous functions in neuronal migration. Development 138, 3033–3042 (2011).

16. Jarjour, A. A. et al. The polarity protein Scribble regulates myelination and remyelination in the central nervous system. PLoS Biol. 13, e1002107 (2015).

17. Moreau, M. M. et al. The planar polarity protein Scribble1 is essential for neuronal plasticity and brain function. J. Neurosci. 30, 9738–9752 (2010).

18. Piguel, N. H. et al. Scribble1/AP2 complex coordinates NMDA receptor endocytic recycling. Cell Rep 9, 712–727 (2014).

19. Hilal, M. L. et al. Activity-Dependent Neuroplasticity Induced by an Enriched Environment Reverses Cognitive Deficits in Scribble Deficient Mouse. Cereb. Cortex 27, 5635–5651 (2017).

20. Robinson, A. et al. Mutations in the planar cell polarity genes CELSR1 and SCRIB are associated with the severe neural tube defect craniorachischisis. Hum. Mutat. 33, 440–447 (2012).

21. Lei, Y. et al. Mutations in planar cell polarity gene SCRIB are associated with spina bifida. PLoS ONE 8, e69262 (2013).

22. Kharfallah, F. et al. Scribble1 plays an important role in the pathogenesis of neural tube defects through its mediating effect of Par-3 and Vangl1/2 localization. Hum. Mol. Genet. 26, 2307–2320 (2017).

23. Wang, L. et al. Digenic variants of planar cell polarity genes in human neural tube defect patients. Mol. Genet. Metab. 124, 94–100 (2018).

24. Murdoch, J. N. et al. Genetic interactions between planar cell polarity genes cause diverse neural tube defects in mice. Dis Model Mech 7, 1153–1163 (2014).

25. Wang, M., Marco, P. de, Capra, V. & Kibar, Z. Update on the Role of the Non-Canonical Wnt/Planar Cell Polarity Pathway in Neural Tube Defects. Cells 8, (2019).

26. Greene, N. D. E. & Copp, A. J. Neural tube defects. Annu. Rev. Neurosci. 37, 221–242 (2014).

27. Copp, A. J. et al. Spina bifida. Nat Rev Dis Primers 1, 15007 (2015).

28. Foss, S., Flanders, T. M., Heuer, G. G. & Schreiber, J. E. Neurobehavioral outcomes in patients with myelomeningocele. Neurosurg Focus 47, E6 (2019).

29. Dauber, A. et al. SCRIB and PUF60 are primary drivers of the multisystemic phenotypes of the 8q24.3 copy-number variant. Am. J. Hum. Genet. 93, 798–811 (2013).

30. Verheij, J. B. G. M. et al. An 8.35 Mb overlapping interstitial deletion of 8q24 in two patients with coloboma, congenital heart defect, limb abnormalities, psychomotor retardation and convulsions. Eur J Med Genet 52, 353–357 (2009).

31. Halevy, A. et al. Microcephaly-thin corpus callosum syndrome maps to 8q23.2-q24.12. Pediatr. Neurol. 46, 363–368 (2012).

32. Stam, A. J., Schothorst, P. F., Vorstman, J. A. & Staal, W. G. The genetic overlap of attention deficit hyperactivity disorder and autistic spectrum disorder. Appl Clin Genet 2, 7–13 (2009).

33. Britanova, O. et al. Satb2 is a postmitotic determinant for upper-layer neuron specification in the neocortex. Neuron 57, 378–392 (2008).

34. Shu, T., Puche, A. C. & Richards, L. J. Development of midline glial populations at the corticoseptal boundary. J. Neurobiol. 57, 81–94 (2003).

35. Pulvers, J. N. et al. Mutations in mouse Aspm (abnormal spindle-like microcephaly associated) cause not only microcephaly but also major defects in the germline. Proc. Natl. Acad. Sci. U.S.A. 107, 16595–16600 (2010).

36. Bishop, K. M., Rubenstein, J. L. R. & O’Leary, D. D. M. Distinct actions of Emx1, Emx2, and Pax6 in regulating the specification of areas in the developing neocortex. J. Neurosci. 22, 7627–7638 (2002).

37. Rachel, R. A., Murdoch, J. N., Beermann, F., Copp, A. J. & Mason, C. A. Retinal axon misrouting at the optic chiasm in mice with neural tube closure defects. Genesis 27, 32–47 (2000).

38. Sun, S. D., Purdy, A. M. & Walsh, G. S. Planar cell polarity genes Frizzled3a, Vangl2, and Scribble are required for spinal commissural axon guidance. BMC Neurosci 17, 83 (2016).

39. Lindwall, C., Fothergill, T. & Richards, L. J. Commissure formation in the mammalian forebrain. Curr. Opin. Neurobiol. 17, 3–14 (2007).

40. Sun, T. et al. A reverse signaling pathway downstream of Sema4A controls cell migration via Scrib. J. Cell Biol. 216, 199–215 (2017).

41. Vaughen, J. & Igaki, T. Slit-Robo Repulsive Signaling Extrudes Tumorigenic Cells from Epithelia. Dev. Cell 39, 683–695 (2016).

42. Gobius, I. et al. Astroglial-Mediated Remodeling of the Interhemispheric Midline Is Required for the Formation of the Corpus Callosum. Cell Rep 17, 735–747 (2016).

43. Yu, H. et al. Frizzled 1 and frizzled 2 genes function in palate, ventricular septum and neural tube closure: general implications for tissue fusion processes. Development 137, 3707–3717 (2010).

44. Yamaguchi, Y. & Miura, M. How to form and close the brain: insight into the mechanism of cranial neural tube closure in mammals. Cell. Mol. Life Sci. 70, 3171–3186 (2013).

45. Pai, Y.-J. et al. Epithelial fusion during neural tube morphogenesis. Birth Defects Res. Part A Clin. Mol. Teratol. 94, 817–823 (2012).

46. Paul, L. K. et al. Agenesis of the corpus callosum: genetic, developmental and functional aspects of connectivity. Nat. Rev. Neurosci. 8, 287–299 (2007).

47. Lu, H.-C. et al. Disruption of the ATXN1-CIC complex causes a spectrum of neurobehavioral phenotypes in mice and humans. Nat. Genet. 49, 527–536 (2017).

48. Nola, S. et al. Scrib regulates PAK activity during the cell migration process. Hum Mol Genet 17, 3552–3565 (2008).

49. Montcouquiol, M. et al. Asymmetric localization of Vangl2 and Fz3 indicate novel mechanisms for planar cell polarity in mammals. J. Neurosci. 26, 5265–5275 (2006).

50. Hong, S.-T. & Mah, W. A Critical Role of GIT1 in Vertebrate and Invertebrate Brain Development. Exp Neurobiol 24, 8–16 (2015).

51. Won, H. et al. GIT1 is associated with ADHD in humans and ADHD-like behaviors in mice. Nat. Med. 17, 566–572 (2011).

52. Dos-Santos Carvalho, S. et al. Vangl2 acts at the interface between actin and N-cadherin to modulate mammalian neuronal outgrowth. Elife 9, (2020).

53. Robert, B. J. A. et al. Vangl2 in the Dentate Network Modulates Pattern Separation and Pattern Completion. Cell Rep 31, 107743 (2020).

54. Moreau, M. M. et al. The planar polarity protein Scribble1 is essential for neuronal plasticity and brain function. J. Neurosci. 30, 9738–9752 (2010).

55. Hilal, M. L. et al. Activity-Dependent Neuroplasticity Induced by an Enriched Environment Reverses Cognitive Deficits in Scribble Deficient Mouse. Cereb. Cortex 27, 5635–5651 (2017).

56. Cervantes-Sandoval, I., Chakraborty, M., MacMullen, C. & Davis, R. L. Scribble Scaffolds a Signalosome for Active Forgetting. Neuron 90, 1230–1242 (2016).

57. El Chehadeh, S. et al. Dominant variants in the splicing factor PUF60 cause a recognizable syndrome with intellectual disability, heart defects and short stature. Eur. J. Hum. Genet. 25, 43–51 (2016).

58. Low, K. J. et al. PUF60 variants cause a syndrome of ID, short stature, microcephaly, coloboma, craniofacial, cardiac, renal and spinal features. Eur. J. Hum. Genet. 25, 552–559 (2017).

59. Santos-Simarro, F. et al. Eye coloboma and complex cardiac malformations belong to the clinical spectrum of PUF60 variants. Clin. Genet. 92, 350–351 (2017).

60. Graziano, C. et al. A de novo PUF60 mutation in a child with a syndromic form of coloboma and persistent fetal vasculature. Ophthalmic Genet. 38, 590–592 (2017).

61. Zhao, J. J. et al. Exome sequencing reveals NAA15 and PUF60 as candidate genes associated with intellectual disability. Am. J. Med. Genet. B Neuropsychiatr. Genet. 177, 10–20 (2018).

62. Moccia, A. et al. Genetic analysis of CHARGE syndrome identifies overlapping molecular biology. Genet. Med. 20, 1022–1029 (2018).

63. Xu, Q. et al. Role of PUF60 gene in Verheij syndrome: a case report of the first Chinese Han patient with a de novo pathogenic variant and review of the literature. BMC Med Genomics 11, 92 (2018).

64. Alkhunaizi, E. & Braverman, N. Clinical characterization of a PUF60 variant in a patient with Dubowitz-like syndrome. Am. J. Med. Genet. A 179, 130–133 (2019).

65. Foss, S., Flanders, T. M., Heuer, G. G. & Schreiber, J. E. Neurobehavioral outcomes in patients with myelomeningocele. Neurosurg Focus 47, E6 (2019).

66. Verheij, J. B. G. M. et al. An 8.35 Mb overlapping interstitial deletion of 8q24 in two patients with coloboma, congenital heart defect, limb abnormalities, psychomotor retardation and convulsions. Eur J Med Genet 52, 353–357 (2009).

67. Dauber, A. et al. SCRIB and PUF60 are primary drivers of the multisystemic phenotypes of the 8q24.3 copy-number variant. Am. J. Hum. Genet. 93, 798–811 (2013).

68. Stergiakouli, E. et al. Shared genetic influences between dimensional ASD and ADHD symptoms during child and adolescent development. Mol Autism 8, 18 (2017).

69. Bush, J. O. & Soriano, P. Ephrin-B1 regulates axon guidance by reverse signaling through a PDZ-dependent mechanism. Genes & Development 23, 1586–1599 (2009).

70. Montcouquiol, M. et al. Asymmetric localization of Vangl2 and Fz3 indicate novel mechanisms for planar cell polarity in mammals. J. Neurosci. 26, 5265–5275 (2006).

71. Pan, C. et al. Shrinkage-mediated imaging of entire organs and organisms using uDISCO. Nature Methods 13, 859–867 (2016).

72. Calderon de Anda, F. et al. Autism spectrum disorder susceptibility gene TAOK2 affects basal dendrite formation in the neocortex. Nature Neuroscience 15, 1022–1031 (2012).

## Supplementary references

1. Madisen, L. et al. A robust and high-throughput Cre reporting and characterization system for the whole mouse brain. Nat. Neurosci. 13, 133–140 (2010).

2. Yamben, I. F. et al. Scrib is required for epithelial cell identity and prevents epithelial to mesenchymal transition in the mouse. Dev. Biol. 384, 41–52 (2013).

3. Hilal, M. L. et al. Activity-Dependent Neuroplasticity Induced by an Enriched Environment Reverses Cognitive Deficits in Scribble Deficient Mouse. Cereb. Cortex 27, 5635–5651 (2017).

4. Ezan, J. et al. Primary cilium migration depends on G-protein signalling control of subapical cytoskeleton. Nature Cell Biology 15, 1107–1115 (2013).

5. Hébert, J. M. & McConnell, S. K. Targeting of cre to the Foxg1 (BF-1) locus mediates loxP recombination in the telencephalon and other developing head structures. Dev. Biol. 222, 296–306 (2000).

6. Gorski, J. A. et al. Cortical excitatory neurons and glia, but not GABAergic neurons, are produced in the Emx1-expressing lineage. J. Neurosci. 22, 6309–6314 (2002).

7. Zhou, L. et al. Early Forebrain Wiring: Genetic Dissection Using Conditional Celsr3 Mutant Mice. Science 320, 946–949 (2008).

8. Montcouquiol, M. et al. Asymmetric localization of Vangl2 and Fz3 indicate novel mechanisms for planar cell polarity in mammals. J. Neurosci. 26, 5265–5275 (2006).

9. Calderon de Anda, F. et al. Autism spectrum disorder susceptibility gene TAOK2 affects basal dendrite formation in the neocortex. Nature Neuroscience 15, 1022–1031 (2012).

10. Moreau, M. M. et al. The planar polarity protein Scribble1 is essential for neuronal plasticity and brain function. J. Neurosci. 30, 9738–9752 (2010).

11. Rivero, O. et al. Cadherin-13, a risk gene for ADHD and comorbid disorders, impacts GABAergic function in hippocampus and cognition. Transl Psychiatry 5, e655 (2015).

